# Multi-objective optimized breeding strategies

**DOI:** 10.1101/209080

**Authors:** Deniz Akdemir, Julio Isidro Sánchez

## Abstract

Multi-objective optimization is an emerging field in mathematical optimization which involves optimization a set of objective functions simultaneously. The purpose of most plant and animal breeding programs is to make decisions that will lead to sustainable genetic gains in more than one traits while controlling the amount of co-ancestry in the breeding population. The decisions at each cycle in a breeding program involve multiple, usually competing, objectives; these complex decisions can be supported by the insights that are gained by using the multi-objective optimization principles in breeding. The discussion here includes the definition of several multi-objective optimized breeding approaches and the comparison of these approaches with the standard multi-trait breeding schemes such as tandem selection, culling and index selection. We have illustrated the newly proposed methods with two empirical data sets and with simulations.

## 1 INTRODUCTION

There are two ways in which the action of a breeder can change the genetic properties of the population; the first by the choice of individuals to be used as parents, which constitutes selection [Falconer et al. (1996); Allard (1999)], and the second by control of the way in which the parents are mated, which embraces inbreeding and crossbreeding [Wright (1921); Kinghorn and Shepherd (1999); Fernández et al. (2001); Sun et al. (2013); Shepherd and Kinghorn (1998); Pryce et al. (2012); Akdemir and Sánchez (2016)]. Selection means breeding from the ”best” individuals whatever ”best” might be. The simplest form of selection is to choose individuals based on their own phenotypic values. Nevertheless, the breeding value is what influences the next generation. If a breeder chooses individuals to be parents according to their phenotypic values, his success in changing the population can be predicted only from knowledge of the degree of correspondence between phenotypic values and breeding values (heritability)[Holland et al. (2003); Cockerham (1963); Dudley and Moll (1969)].

Breeders have been selecting on the basis of phenotypic values since domestication of plants and animals or, more recently, breeders have substantially used the pedigree-based prediction of genetic values for the genetic improvement of complex traits Henderson (1984); Gianola and Fernando (1986); Crossa et al. (2006); Piepho et al. (2008).

Since the invention of the polymerase chain reaction by Mullis in 1983, the enhancements in high throughput genotyping [Lander et al. (2001); Margulies et al. (2005); Metzker (2010)] have transformed breeding pipelines through marker-assisted selection (MAS) [Lande and Thompson (1990)], marker assisted introgression [Charcosset and Hospital (1997)], marker assisted recurrent selection [Bernardo and Charcosset (2006)], and genomic selection (GS) Meuwissen et al. (2001). The latter use genome-wide markers to estimate the effects of all genes or chromosome positions simultaneously [Meuwissen et al. (2001)] to calculate genomic estimated breeding values (GEBVs), which are used for selection of individuals. This process involves the use of phenotypic and genotypic data to build prediction models that would be used to estimate GEBV’s from genome wide marker data. It has been proposed that GS increases the genetic gains by reducing the generation intervals and also by increasing the accuracy of estimated breeding values. However, many factors are involved in the relative per unit of time efficiency of GS and its short and long time performance [Jannink et al. (2010); Daetwyler et al. (2007)].

When selection is applied in a breeding program to the improvement of the economic value of an individual, it's generally applied to several traits simultaneously and not just to one, because economic value depends on more than one trait [Lynch et al. (1998); Bernardo (2002)]. Hence deciding which are the most valuable individuals to select for parents of the next generation forces the breeder to consider several different characteristics. This is usually referred to as multiple trait selection and implied selection for correlated traits. These are not likely all to be equally important or all to be independent of each other. Correlated traits are of interest for two main reasons in breeding programs. Firstly, to understand the genetic causes of correlation through the pleiotropic action of genes. Secondly, because it is key to understand how the improvement of one trait will trigger concurrent changes in other traits [Allard (1999)).

For example, the genetic improvement of grain yield (GY) and grain protein concentration (GPC) is impeded by several factors. Firstly, increasing both GY and GPC represents a major challenge in plant breeding due to the observed negative correlation [Cooper et al. (2001); Oury and Godin (2007); Zheng et al. (2009); Groos et al. (2003); Kibite and Evans (1984); Rharrabti et al. (2001); DePauw et al. (2007); Simmonds (1995)). Secondly, future GY improvements must be achieved under global warming, with a reduction of the use of water and fertilizers due to environmental issues. At the same time, GPC must be at least maintained at its current level, since grain protein is the major source of dietary proteins for humans [Shewry (2007); Aiking (2011). Therefore, selecting multiple plant traits associated with crop yield improvement is difficult, especially if consider the genotype x environment (G × E) interactions that lead to complicate selection for broad adaptation [de la Vega (2012); Cooper and DeLacy (1994)] and the low heritability of traits [Moose and Mumm (2008)].

There are many ways of selecting for several different traits but these will not often be equally efficient. The most efficient method is that which results in the maximum genetic improvement per unit of time and effort expended [Smith (1936); Hazel and Lush (1942)].

One might select in turn of each trait singly in successive generations (tandem selection) until each trait is improved to a desired level. Tandem selection is practical when some traits can be meaningfully evaluated in the earlier stages of a breeding program and other traits can be evaluated only later [Hallauer and Miranda (1987); Acquaah (2009); Burgess and West (1993)]. One might select for all the traits at the same time but independently, rejecting all individuals that fail to come up to a certain standard for each trait regardless of their values for any other of the traits (independent culling levels) [Hazel (1943)]. Only individuals that meet the minimum or maximum standards for each trait are selected. Nevertheless, the method that has shown the most rapid improvement of economic value, is to apply the selection simultaneously to all the component traits together, appropriate weight being assigned to each trait according to its relative economic importance, its heritability and the genetic and phenotypic correlations between the different traits [Hazel and Lush (1942); Hazel (1943); Williams (1962)]. The component traits should be combined into a score or index, in such a way that selection applied to the index, as if the index were a single trait, will yield the most rapid possible improvement of economic value.

The multitrait breeding problem arise a fundamental question in terms of the best procedure to reach the breeding goals. In recent years, powerful numerical and statistical methods and tools have been developed to model uncertainty and multitrait selection in plant breeding not just to improve selection [Meuwissen et al. (2001); Goddard (2009); Bernardo and Yu (2007); Heffner et al. (2009)] but also as a tool to facilitate ideotype design in crop modelling [Picheny et al. (2017); Gouache et al. (2017); Casadebaig et al. (2016); Martre et al. (2015b)]. Numerical models can predict the outcome of plant traits by simulating physiological processes and their interaction with the environment [Rötter et al. (2015); Ghanem et al. (2015); Martre et al. (2015a)]. Nevertheless, there is a need of approaches that explorer deeper the optimal trait combinations.

In this article, we proposed an approached based on a multi-optimization framework by setting optimal compromise solutions (Pareto front) that should be identified by an effective and complete search procedure to let the breeder to carry out the best choice. Our approach extends and generalizes the previously proposed approaches mentioned above and optimized parental contribution for single and multiple trait breeding. The aim objectives of this article are i) set up a multi-optimization framework for plant breeding and ii) to compare the efficiency of our method with previous multiple trait approaches.

## 2 MATERIALS

### 2.1 Wheat and barley datasets

The genetic material used in this study consists of two different datasets on wheat and barley. Both of these data sets were downloaded from the triticeae toolbox (https://triticeaetoolbox.org) and more details about these datasets are provided in Table 1.

**TABLE 1.**
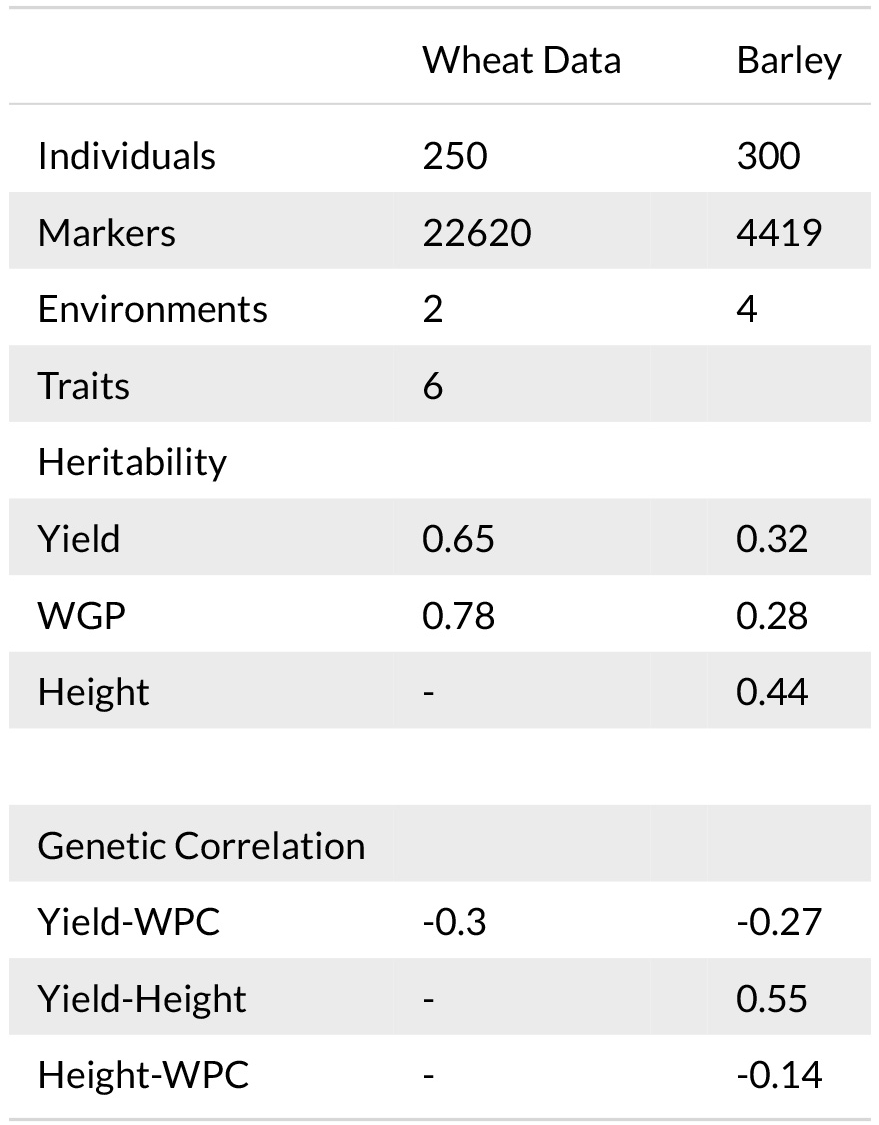
Wheat and Barley Datasets

Grain protein content is an important trait for the nutritional value of grain. Grain protein content in wheat, barley and oats is usually negatively correlated with grain yield, thus increase in protein quantity often results in yield reduction. We will use a wheat and a barley data to illustrate the use of multi-objective optimized breeding approaches that were suggested in the previous sections.

#### Model for estimating breeding values

For each dataset, the genomically estimated breeding values (GEBVs) for the traits yields and grain protein were predicted from a multi-trait mixed model using the accounting for environmental effects as fixed effects and breeding values of individuals as a random effect coming from a zero centered matrix-variate normal distribution with a seperable (kronecker product) covariance structure for traits and genotypes. A similar model was assumed for the random error terms. More specifically, the model that is used is given by

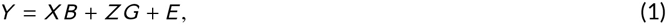

where *Y* is the *n* × *d* response variable, *X* is the *n* × *q* design matrix of *q* × *d* the fixed effects *B*, *Z* is a *n* × *q*_*j*_ design matrix of the *q* × *d* random effects *G* and *E* is the *n* × *d* matrix of residual effects. The random effects and the residual are independently distributed, and have matrix variate distributions (*G* ~ *N*_*q*×*d*_(0_*q*×*q*_, *K*, ∑) and (*E* ~ *N*_*n*×*d*_(0_*n*×*d*_, *K*, ∑_*E*_)). The GEBVs obtained from the above model were standardized by dividing each with their corresponding standard deviations to to bring the GEBVs to the same scale.

### 2.2 Simulations

The long term performance was evaluate by simulations. Beginning with two founders we have formed a population of N (N=100, 200, 300, 400) genotypes with 1000 single nuceotide polymorphism (SNPs) at 3 chromosomes each and carried this population through 100 generations of random mating. Two traits were defined simultaneously by attaching random quantiatative trait loci (QTL) effetcs at 200 randomly selected loci on each chromosome where 100 of these were taken to have opposite sign effects for the different traits, using these effects to calculate the genotypic values for each individual and adding to each of these a value generated from a normal distribution with zero mean and variance equal to the variance of the genotypic values in the population. The base population for breeding simulations were obtained by simulating 10 rounds of tandem selection with 50% selection intensity. Starting from this base population, the results from 30 replications of 16 rounds of GS and 10 rounds of PS with tandem, index selection and culling and 16 rounds with three multi-trait breeding methods recommended in this manuscript have been simulated. Marker effects were estimated from phenotypic and genotypic data on the current population data at odd numbered breeding cycles.

## 3 METHODS

In this section we will describe two new approaches along with the standard methods for multi-trait breeding. We will illustrate and compare these methods with two empirical data sets and with simulation studies. More details about the subject can be found in Deb (2001); Konak et al. (2006); Coello (2006) and references there in.

### 3.1 Multi-objective optimization and related concepts

A single-objective optimization problem is defined as the minimization (or maximization) of a scalar objective function *f*(*x*) subject to inequality constraints *g*_*i*_(*x*) ≤ 0, *i* = 1,…, *m* and equality constraints *h*_*j*_(*x*) = 0, *j* = 1,…,*p* where *x* is a *n*-dimensional decision variable vector.

Real-world problems may require a simultaneous optimization of multiple objectives. In problems with a single goal (mono-objective), optimization algorithms look for the best solution in the search space, or in a set of solutions. On the other hand, in multi-objective problems, there may not be an optimal solution for all objectives. In this case, in a multi-objective problem there exists a subset in the search space which is better than the rest of the solution. This subset is known as Pareto optimal solutions or non-dominated solutions.

More formally, multi-objective problems are those problems where the goal is to optimize simultaneously *k* objective functions designated as: *f*_*1*_(*x*), *f*_*2*_(*x*),…,*f*_*k*_(*x*) and forming a vector function 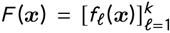 subject to inequality constraints *g*_*i*_(*x*) ≤ 0,*i* = 1,…, *m* and equality constraints *h*_*j*_(*x*) = 0, *j* = 1,…,*p*.

Although single-objective optimization problems may have a unique optimal solution, multi-objective problems (as a rule) present a possibly uncountable set of solutions. In order to let the decision maker, the designer, to carryout the best choice the set of optimal compromise solutions (Pareto front) has to be identified.

Pareto optimal solutions are those solutions within the decision space whose corresponding variables cannot be all simultaneously improved. These solutions are also termed non-inferior, admissible or efficient solutions. Their corresponding vectors are termed non dominated. In multi-objective optimization problem, the goodness of a solution is determined by the dominance. By selecting a vector from this vector set implicitly indicates acceptable Pareto optimal solutions, decision variables. These solutions may have no apparent relationship besides their membership in the Pareto optimal set. They form the set of all solutions whose associated vectors are non dominated.

A vector *u* = (*u*_1,…,*u*_ *k*) is said to dominate another vector *v* = (*v*_1_,…,*v*_*k*_) (written as *u* ⪰ *v*) if and only if *u* is partially less than *v*,i.e., ∀*i* ∈ 1,…,*k*, *u*_*i*_ ≤ *v*_*i*_ Λ ∃*i* ∈1,…,*k u*_*i*_ < *v*_*i*_. Pareto optimal solutions are those which, when evaluated, produce vectors whose performance *f*_*i*_ cannot be improved without adversely affecting another *f*_*j*_, *i* ≠ *j*. In a minimization problem, a solution *x* is said to be Pareto optimal if and only if there is no *x′* for which *F*(*x′*) dominates *F*(*x*), i.e., there exists no feasible vector *x′* which would decrease some criterion without causing a simultaneous increase in at least one other criterion.

For a multi-objective problem, *F*(*x*), the Pareto Optimal Set, *P*\*, is defined as: *P*\*:= {*x*: ¬∃*x F*(*x′*) ⪰ *F*(*x*)}.

The Pareto front *PF*\* is defined as: *PF*\*:={*u* = *F*(*x*)|*x* ∈ *P*\*}.

Non-dominated ordering implements the concept of dominance and classifies a population of solutions into boundaries according to their level of dominance. The first level includes all the non-dominated solutions, the second level are formed by the non-dominated solutions after excluding the solutions in the first level and this allocation process finishes when all solutions are allocated within their respective frontiers. After the this process, the first-frontier solutions are not dominated by any other individual; however, they dominate the second frontier. Thus, solutions of the *i*th frontier dominate individuals of the (*i* + 1)th frontier, i.e, solutions can be sorted according to these frontiers.

#### Selecting a “good” solution on the frontier surface

At the end of a multi-objective optimization, the decision maker (DM) has to select the preferred solutions from the Pareto frontier; this can be a difficult task for high dimensional multi-objective optimization problems. For this reason, decision making support tools are developed to aid the DM in selecting the preferred solutions. The choice of a unique solution in the collection of Pareto optimal solutions depends on the knowledge of problem characteristics, and a solution in a particular model may not be the best in another model or environment. For 2 and 3 dimensional multiobjective optimization problems a strategy is to first plot the Pareto frontier followed by visual identification of the kink (leg) of the frontier as the region of preferred solutions. For more than three objective problems, selecting the preferred solutions becomes a difficult issue since there is no easy or intuitive method to visually represent the Pareto frontier. A few decision support tools were described in Zio and Bazzo (2012); Agrawal et al. (2005), and Tušar and Filipič (2015).

An heuristic approach for identifying preferred solutions on the frontier can be defined by using the ideal solution concept and global criterion: Let 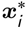 be a vector which optimizes the *i*th objective function *f*_*i*_(*x*) for *i* = 1,2,…,*k* Then the vector 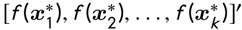 is ideal for an multi-objective problem and is consequently called the ideal vector. The global criterion method aims to minimize a function (global criterion) which is a measure of how close the decision maker can get to the ideal vector. A measure of closeness to the ideal solution is a family of *L*_*p*_-metrics defined as follows:

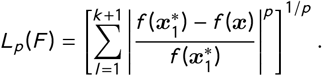

If the functions values are normalized to the range [0,1], then the above formula becomes

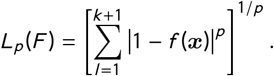

Finally, if weights are attached to the functions *f*_*1*_(*x*), *f*_*2*_(*x*),…,*f*_*k*+1_(*x*) a weighted version can be written as

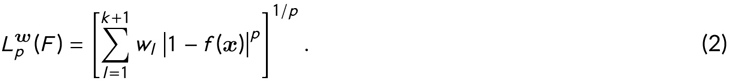

In the remaining of this manuscript, we have used *p* = 2 which coincides with the Euclidean distance.

#### Self-Organizing Maps for visualizing the Pareto optimal solutions

Self-Organizing Maps (SOMs) [Kohonen (1981, 1998)] have been recommended for visualizing the pareto optimal solutions for high dimensional multi-objective problems [Obayashi and Sasaki (2003)]. Neural networks are used in learning tasks that are too complex for human brain to comprehend and SOM is a unsupervised neural networks technique for organizing complex or vast amounts of data by providing lower dimensional representations of data in manner that is most easily understood. Specifically, SOMs are a type of artificial neural network (ANN) that provides a topology preserving mapping from the high dimensional space to map units. The property of topology preserving means that the mapping preserves the relative distance between the points; points that are near each other in the input space are mapped to nearby map units in the SOM. The SOM can thus serve as a cluster analyzing tool of high-dimensional data and be used as a visual aid in determining a ’good’ solution on the frontier surface.

### 3.2 Multi-objective optimized breeding strategies

Genomic selection (GS) is being used increasingly in plant breeding to accelerate genetic gain (Crossa et al. 2010; Roorkiwal et al. 2016; Zhang et al. 2015; Edriss et al. 2017). Genomic selection focuses on best performance of parents before mating, while genomic mating (GM) (Akdemir 2017) includes information on complementation of parents to be mated and thereby is more sustainable in the longer term.

Some standard ways of breeding for gains in multiple selection include tandem selection, culling and index selection. In tandem selection, the breeder aims to improve only one of the traits at each breeding cycle while sequentially alternating among the traits through the breeding cycles. Culling involves removal of inferior individuals from the breeding population one trait at a time starting with a very low intensity of selection and sequentially alternating over the traits; following this a second, third, … rounds of elimination are performed by increasing the selection intensity slightly at each round until a preferred number of individuals are left in the population. These individuals are mated to create the next generation of breeding population. Index selection involves the breeder to assign economic weights to each trait and selection is done using the weighted scores of individuals. The standard breeding approaches, such as PS, GS, GM and pedigree-based prediction, can be used with any of these multi-trait breeding approaches.

The selection index has been shown to be the most efficient method for maximum aggregate genetic progress, provided that(1) reliable estimates of genotypic and phenotypic variances and covariances are available and (2) appropriate economic weights of each trait can be determined. Nevertheless, economic weights are not easily determined (Pesek and Baker (1969)) and do not represent the genetic potential in the breeding population.

In the remaining of this article, we assume that a high density marker data is available for the current breeding population from which the co-ancestry coefficients can be calculated, and that there is no pedigree information. In view of the reducing genotyping costs, mostly incomplete or non-existing pedigrees in plant populations, this assumption is a reasonable one. The implementation of PS in our simulations did not use any genotypic information or pedigrees. Basically, it referred to selecting the individuals with best observed phenotypes to be parents in the next generation. Results elsewhere (Forni et al., 2011) indicate that there would be no significant differences between PS and GS if a pedigree from many generations is used in pedigree based estimation of BV’s. In addition, there are methods to combine pedigrees with marker based relationship matrices (Legarra et al., 2009; Meuwissen et al., 2011) which would result in a yet another selection approach.

#### Non-dominated selection

The first type of multi-trait breeding approach we discuss makes use of the non-dominated sorting concept. One approach involves sorting the individuals in a breeding program according to non-dominance ordering using their (predicted) breeding values over the traits of interest. The assignments of parental contributions are done by assigning higher weights to individuals at lower non-dominance order, and this approach could also be coupled with other selection strategies as elitism.

A similar approach involves enumeration of all possible subsets of the traits and counting the number of times each individual is non-dominated over these subsets. Thus is followed by assigning higher parental contribution proportions to individuals with higher counts.

#### Multi-objective optimized genetic gains while controlling co-ancestry

Let *A* be a matrix of pedigree based numerator relationships or the additive genetic relationships between the individuals in the genetic pool (this matrix can be obtained from a pedigree of genome-wide markers for the individuals) and let *c* be the vector of proportional contributions of individuals to the next generation under a random mating scheme. The average inbreeding and co-ancestry for a choice of *c* can be defined as *r* = 1/2*c′Ac*. If *b* is the vector of GEBV’s, i.e., the vector of BLUP estimated BV’s of the candidates for selection. The expected gain is defined as *g*(*c*) = *c′b*. Without loss of generality, assume that the breeder’s long term goal is to increase the value of *g*(*c*).

Several authors [Wray and Goddard (1994); Brisbane and Gibson (1995); Meuwissen (1997); Meuwissen et al. (2001); Sonesson et al. (2012); Clark et al. (2013)] have proposed minimizing the average inbreeding and co-ancestry while restricting the genetic gain. These approaches find the parental proportions obtained by solving the following optimization problem: minimize *r*(*c*) = 1/2*c′Ac* subject to *c′b* = *ρ*, and *c′*1 = **1***′c* ≥ 0, where *ρ* is the desired level of gain. This problem is easily recognized as a quadratic optimization problem (QP). There are many efficient algorithms that solves QP’s so there is in practice little difficulty in calculating the optimal solution for any particular data set. Recently, several allocation strategies were tested using QP’s in Goddard (2009); Pryce et al. (2012); Schierenbeck et al. (2011).

We suggest the following extension of the above formulation for obtaining parental proportions in the multi-trait scenario: Let *g*_1_, *g*_2_,…,*g*_*k*_ denote the *m*-dimensional vectors of GEBVs for *k* traits. Without loss of generality assume a maximum is sought for all these traits. As in Wray and Goddard (1994); Brisbane and Gibson (1995); Meuwissen (1997); Meuwissen et al. (2001) we want to keep inbreeding to minimum. This defines the multi-objective optimization problem; more formally we are looking to solve the maximization of the vector function

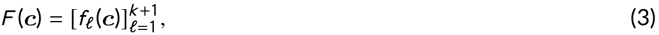

with 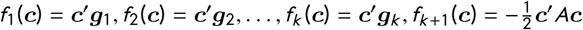 subject to inequality constraints *c*_*i*_ ≥ 0, *i* = 1,…,*m* and equality constraint 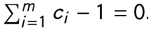. This formulation defines a QP.

A slight modification of the above aims to penalize negative genetic correlations between trait pairs, and it involves changing 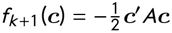 to

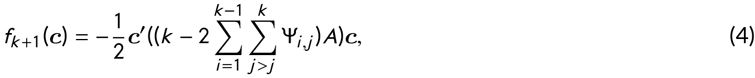

where ψ_*i,j*_ denotes the genetic correlation between traits.

The second class of multi-trait breeding schemes assigns parental proportions using one of the preferred solutions on the Pareto surface for the multi-objective problems stated in Equations (3) or (4).

Since each of the components of these two optimization problems are concave, Kuhn-Tucker conditions (KTC, see Supplementary file for more details) are not only necessary but also sufficient for a solution to be Pareto optimal, therefore, the whole set of Pareto optimal solutions are characterized as the solutions for all possible choices of 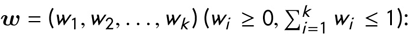:

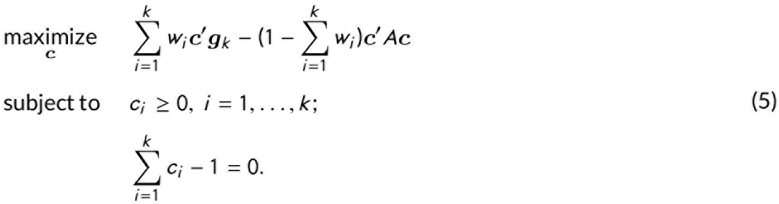

The term *c′Ac* would be need to be changed with 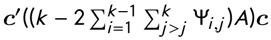 for the second formulation.

##### Ideal Solution

A simple estimator, say *maxf*_*i*_, for the ideal solution 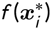, for trait *i* in a certain breeding population is the maximum observed value for that trait. It is also possible to estimate this quantity by calculating the maximal estimated genomic value using the marker effects estimates. We use the former approach in the remaining of this article. When calculating the distance from the ideal solution, the objective function values were scaled to the range [0, 1] using the transformation

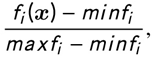

for trait *i*; and *minf*_*i*_ is the estimate for the worst value of the trait *i*, it is calculated in the same fashion as *maxf*_*i*_.

#### Genomic mating in the context of multi-objective optimized breeding

As opposed to the continuous parentage contribution proportions solutions in the GS method, the genomic mating (GM) method gives the list of parent mates of the progeny. Multi-objective optimization problem (assuming maximization is sought for the trait) of the GM problem involves minimization of *−Gain*(*P*), *−Usef ulness*(*P*) and *Inbreeding*(*P*) with respect to mating plan *P*.

The expected gain for a mating plan can be calculated from the mid parent genetic values. There are several alternative measures of inbreeding based on mating plans Leutenegger et al. (2003); Wang (2011). In Akdemir and Sánchez (2016), we have used a measure derived under the infinitesimal genetic effects assumption proposed by Quaas (1988) and Legarra et al. (2009). Measures of expected cross-variance usefulness) can be obtained using the results in Akdemir and Sánchez (2016) under the assumption of unlinked markers. An alternative approach would be to use simulated progenies to calculate the cross-variances. One can easily include information about the LD in these simulations. Yet another measure of usefulness was proposed in Zhong and Jannink (2007).

Extention to multi-trait genomic mating for a *k* trait problem (assuming maximization is sought for traits) is defined by the optimization problem which seeks minimization of *−Gain*(*P*)_*i*_, *−Usefulness*(*P*)_*i*_, for *i* = 1 :, 2,…, *k* and *Inbreeding*(*P*) with respect to mating plan *P*.

## 4 RESULTS

### 4.1 Wheat and barley data sets

Figure 1 and Supplementary Fig. S3 show the non-dominated selection solutions for wheat and barley datasets. The total number of individuals for each dataset are represented with different colors depending on the non-dominated level. These solutions within the decision space from the Pareto optimal correspond to variables that cannot be all simultaneously improved. That is to say, you cannot improve both grain protein and yield when selecting one line (circle) within a non-dominated level. The most optimal solution should be found on the right top corner of the graph but this is not feasible. Therefore, a breeder should select lines balancing both traits simultaneously.

**FIGURE 1.**
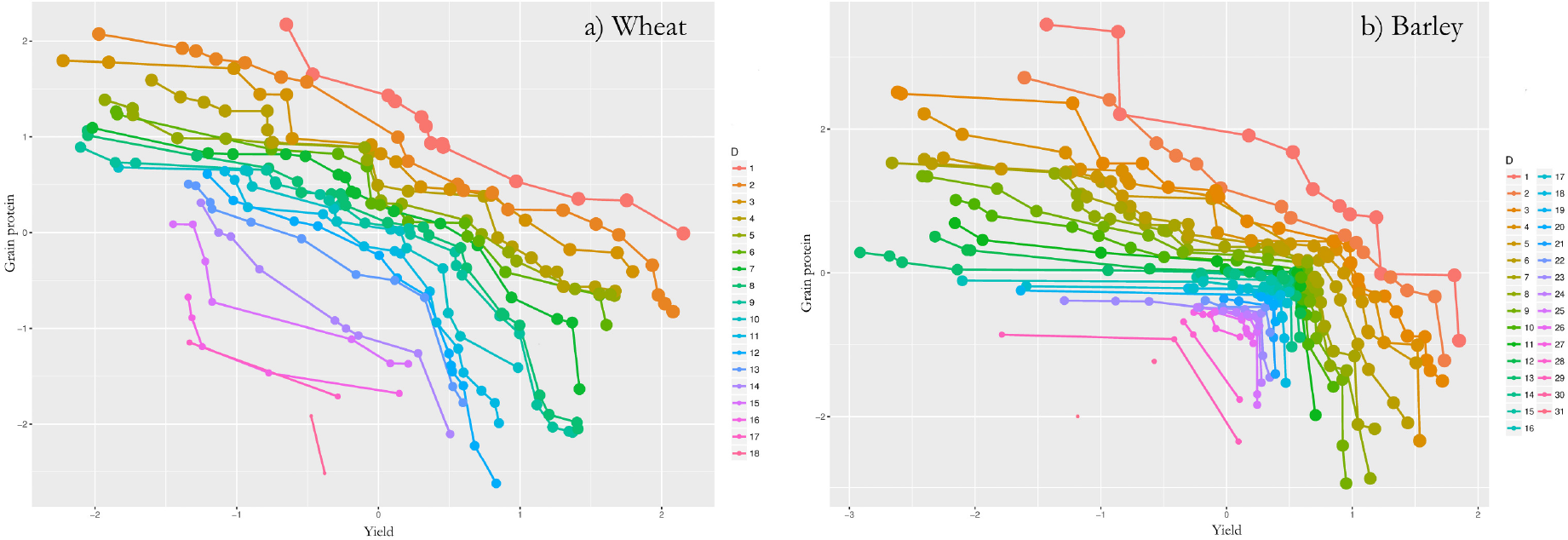
These figures are obtained by plotting GEBVS from model in (1) for grain yield and grain protein content in wheat (a) and barley (b) dataset. The point and lines with the same color belong to the same dominance level. The dominance level 1 is obtained by identifying all of the non-dominanted individuals. Level 2 is obtained as the non-dominated individuals in this smaller subset obtained by removing the individuals in Level 1. This process is continued until all the indivudals are assigned to their dominance levels. There are 18 and 33 levels of dominance for wheat and barley data sets.

The Supplementary Fig. S3 shows the non-dominated solutions on three dimensional space.

The frontier surface related to the optimal parental proportions for the wheat and barley datasets are given in Figures 2 (a) and 2 (b). These figures show the frontier surfaces obtained by plotting Pareto optimal solutions for parental contributions obtained by solving the optimization problem given in Equation (3) for improving yield and protein content while controlling coancestory. The redness of the points indicates closeness to ideal solutions as calculated by Equation (2).

**FIGURE 2.**
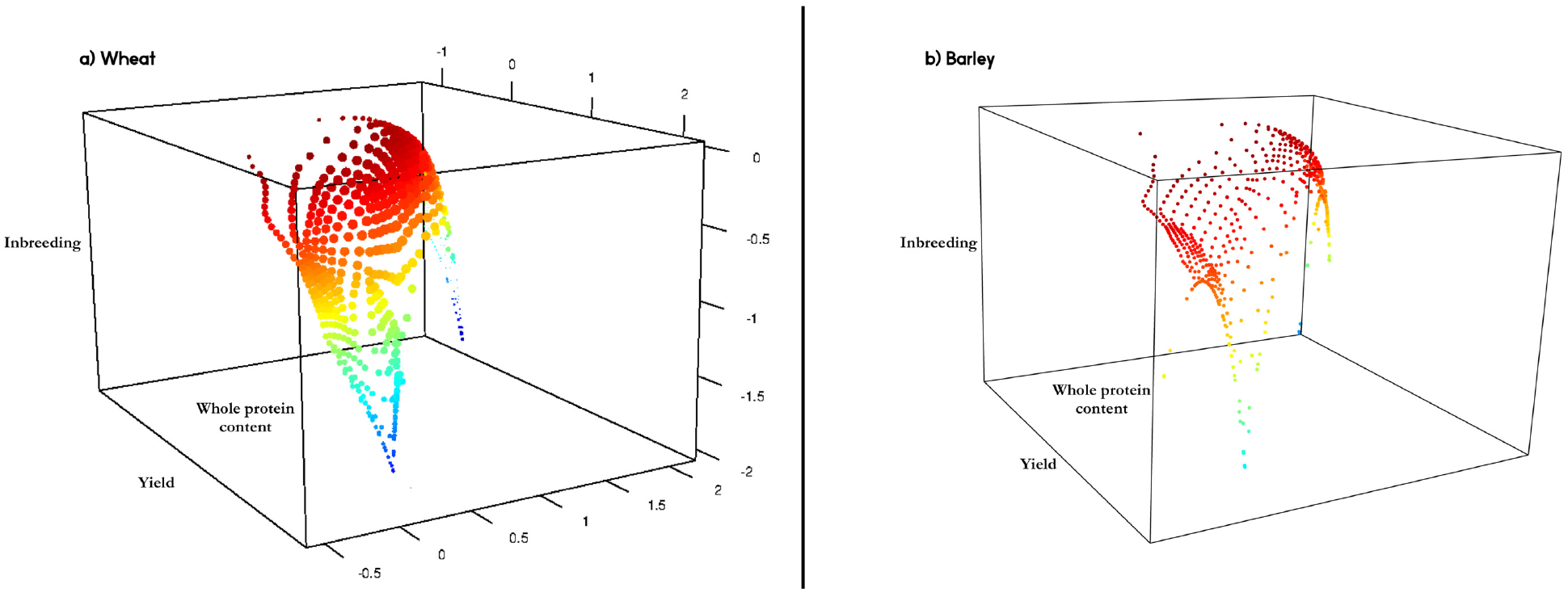
These figures show the frontier surfaces obtained by plotting Pareto optimal solutions for parental contributions obtained by solving the optimization problem giving in Equation (3) for improving yield and protein content while controlling coancestory. The redness of the points indicates closeness to ideal solutions as calculated by Equation (2).

Figure 3 and Supplementary Figure S5 show an example of three "good" solutions on the wheat and barley frontier curve obtained from Figure 2 as calculated by Equation (2), i.e., the solutions on the frontier curve were ranked according to their distance from the ideal solution and these figures correspond to the parental contribution solutions that are closest to the ideal. Note that non-dominated ordering based approaches (1) give similar solutions to optimized parental proportions approach. Nevertheless, solutions for the parental proportions represent the control in inbreeding and have different weights for the individuals on the same level of dominance.

**FIGURE 3.**
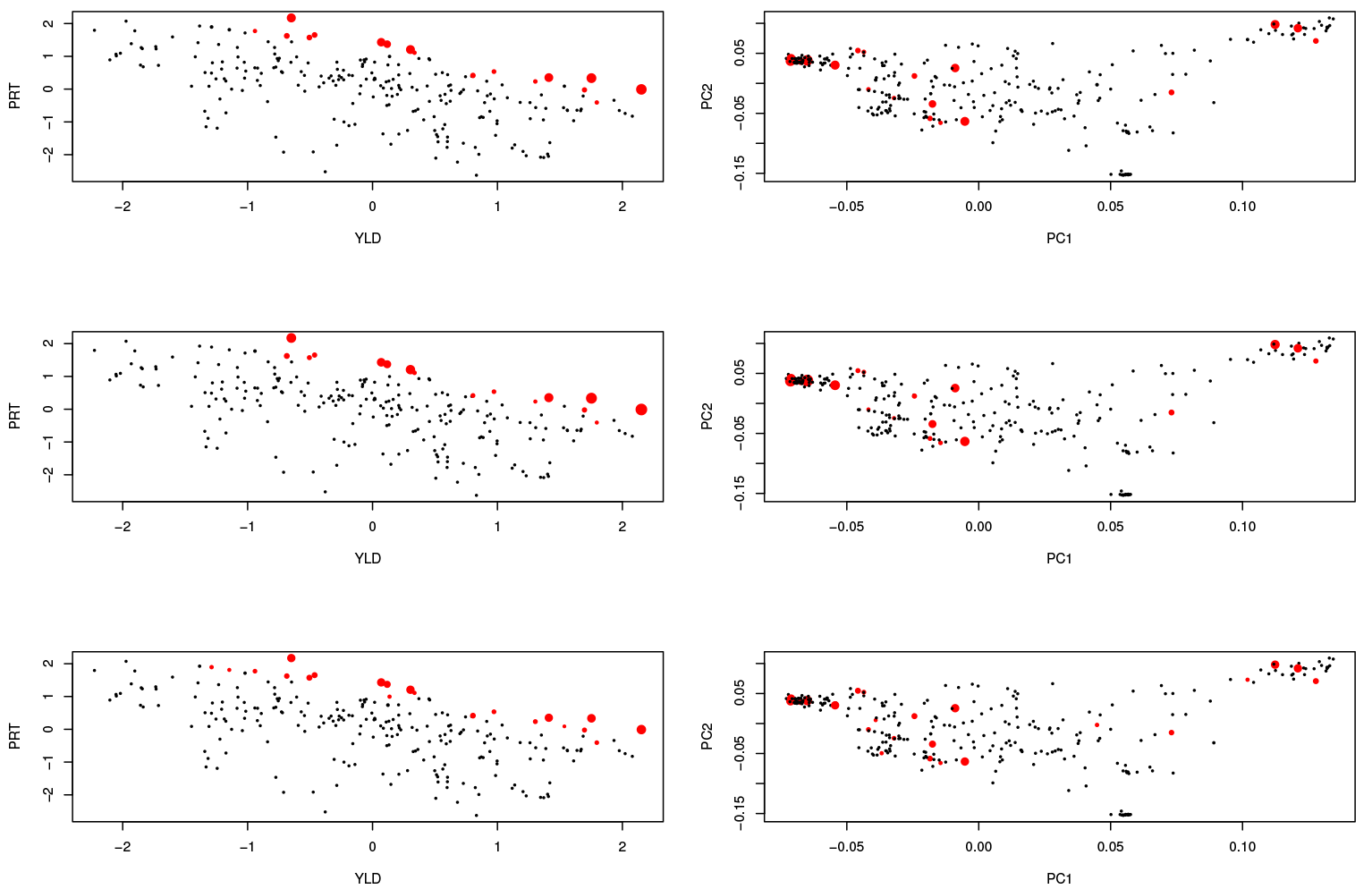
Three ’good’ solutions on the wheat frontier curve obtained from Figure 2. Red points indicated the individuals that have non-zero parental proportions. The size of the points are proportional to the magnitude of the parental contributions. The figures on the right side, represent the same information but on the first two principal components of the genotyping marker space. YLD: Yield; PRT: Protein; PC: Principal components

The multi-trait breeding described in this manuscript can also be used to improve one or more traits in several target environments. For example, a common breeding goal is improvement of yield in multiple environments. Barley data was collected in 4 environments. The non-dominance ordering for the GEBVs for yield in four environments (dry-irrigated x high-low nitrogen) are given in Figure 6.

**FIGURE 4.**
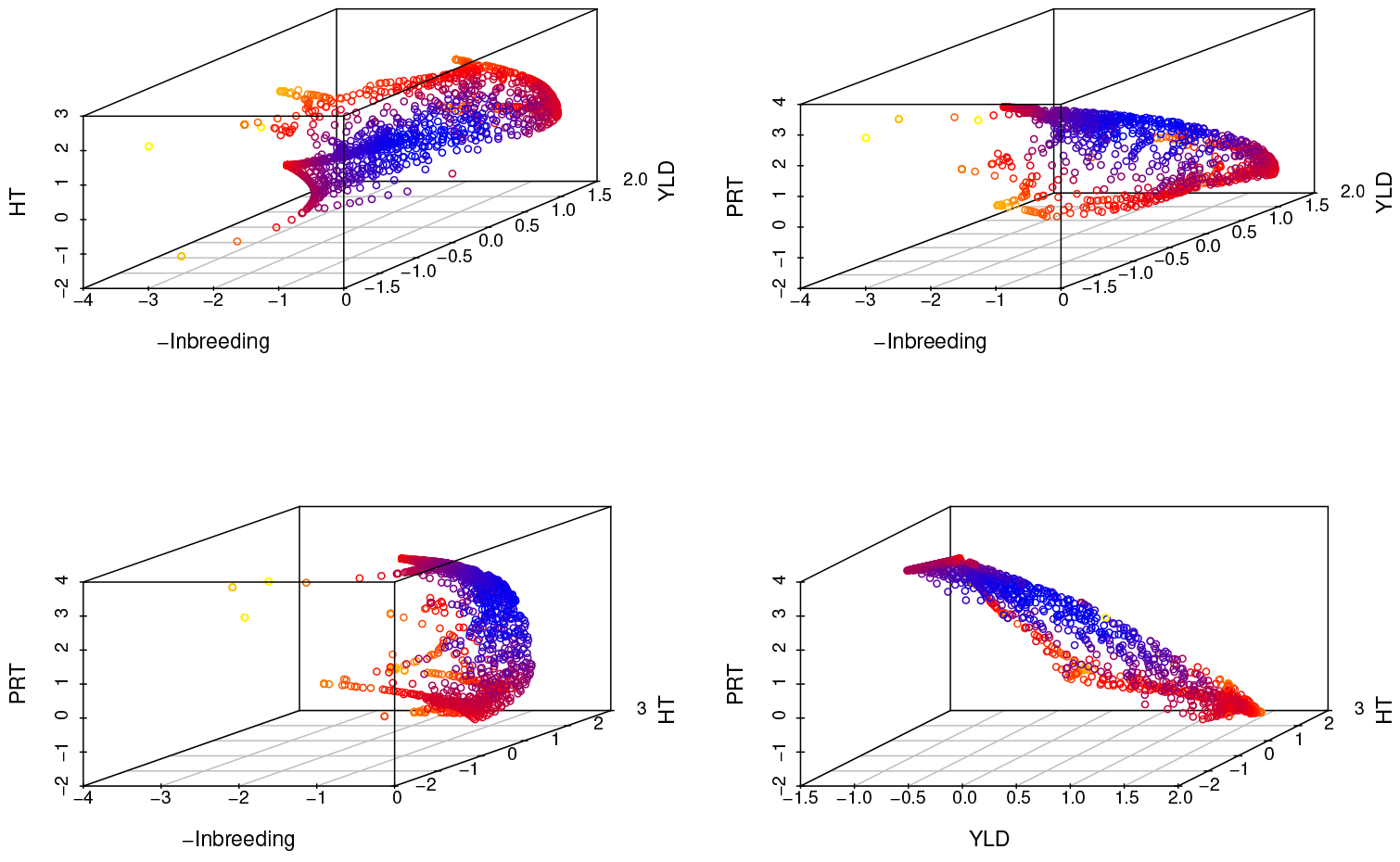
Barley data: The dimensions correspond to negative of inbreeding, and average gains based on GEBVs for yield, height and protein content. Each point on the 3 dimensional scatterplots correspond to a Pareto optimal solution for parental contributions. Blueness of the points measure the closeness to the ideal solution as calculated by Equation (2). HT: Height; PRT: Protein; YLD: Yield

**FIGURE 5.**
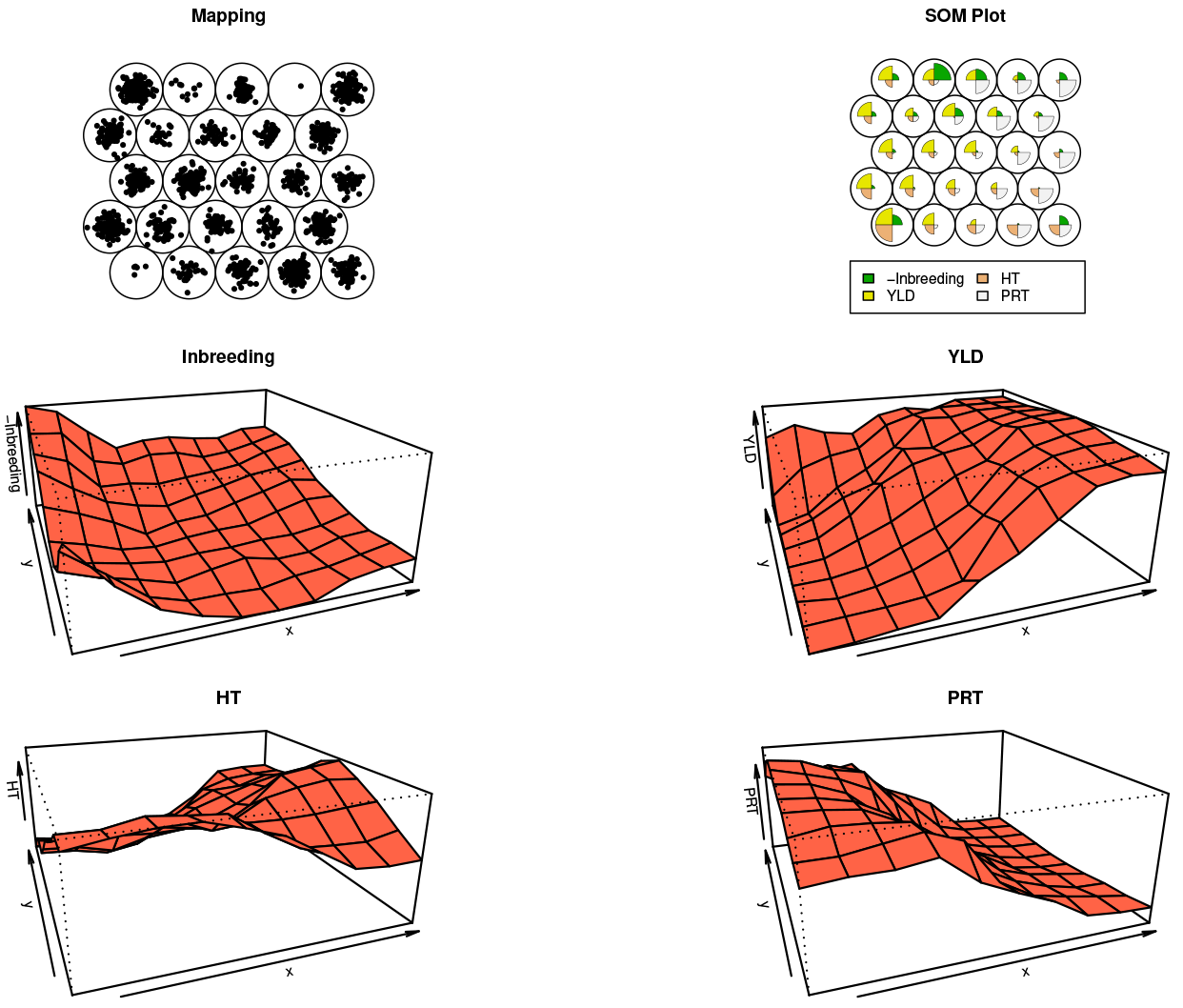
Self-Organizing Maps plot for barley data for parental proportions over three traits, yield, protein content and height and negative of inbreeding. This is another representation of the fourth dimensional Pareto surface in Figure 4.

**FIGURE 6.**
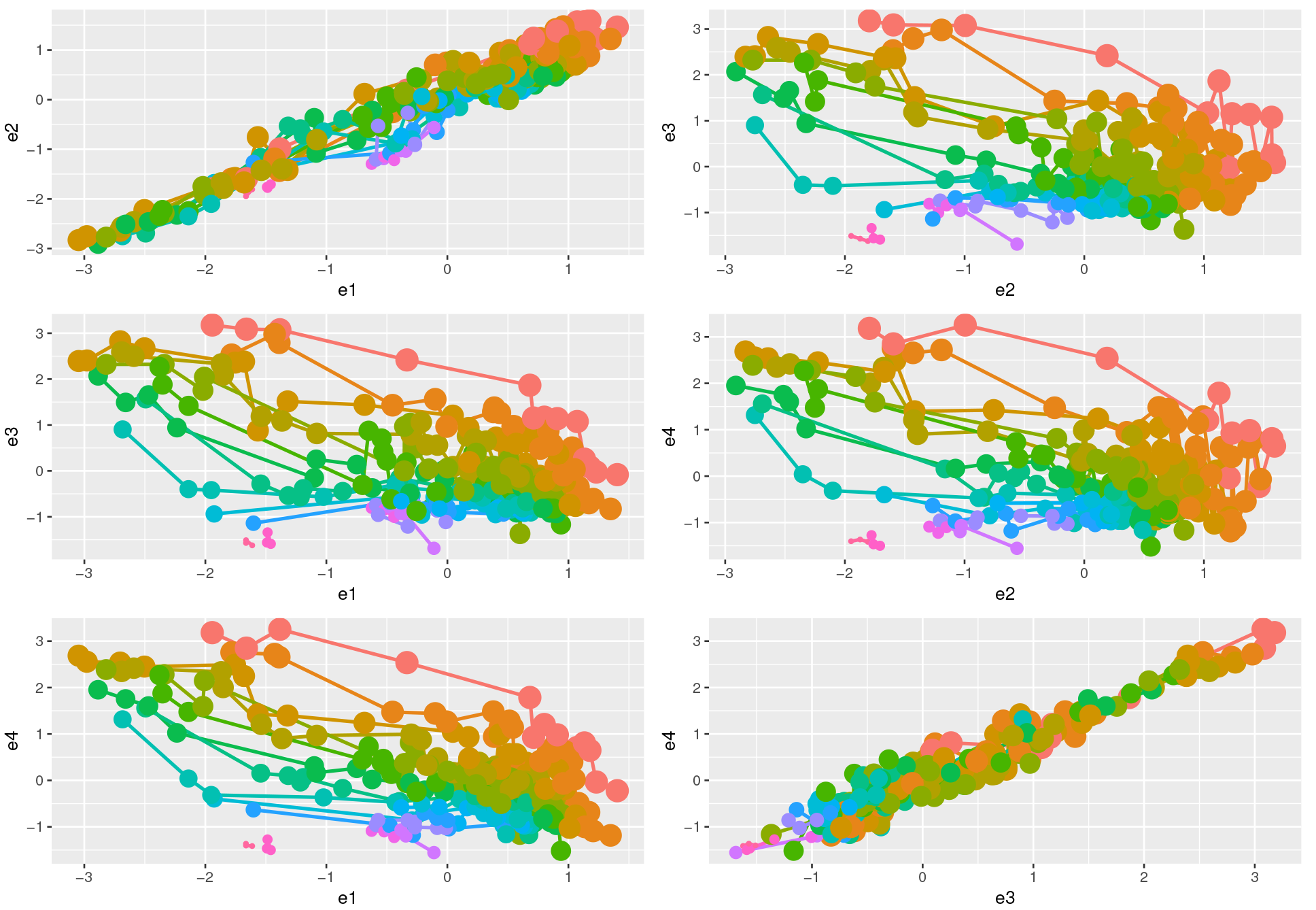
Barley data: Dominance ordering based on yield in 4 environments (dry-irrigated × high-low nitrogen). The environment specific GEBVs for grain yield for barley data are plotted with the dominance ordering of these individuals indicated by the lines of different color.

### 4.2 Longterm performance evaluated by simulations

Figure 7 (a-d) show the results from simulations for the study of the long term behavior of PS, GS using the standard methods Index and tandem selection and culling along with the results for nondominated selection, and the two forms of multi-objective optimized breeding schemes.

**FIGURE 7.**
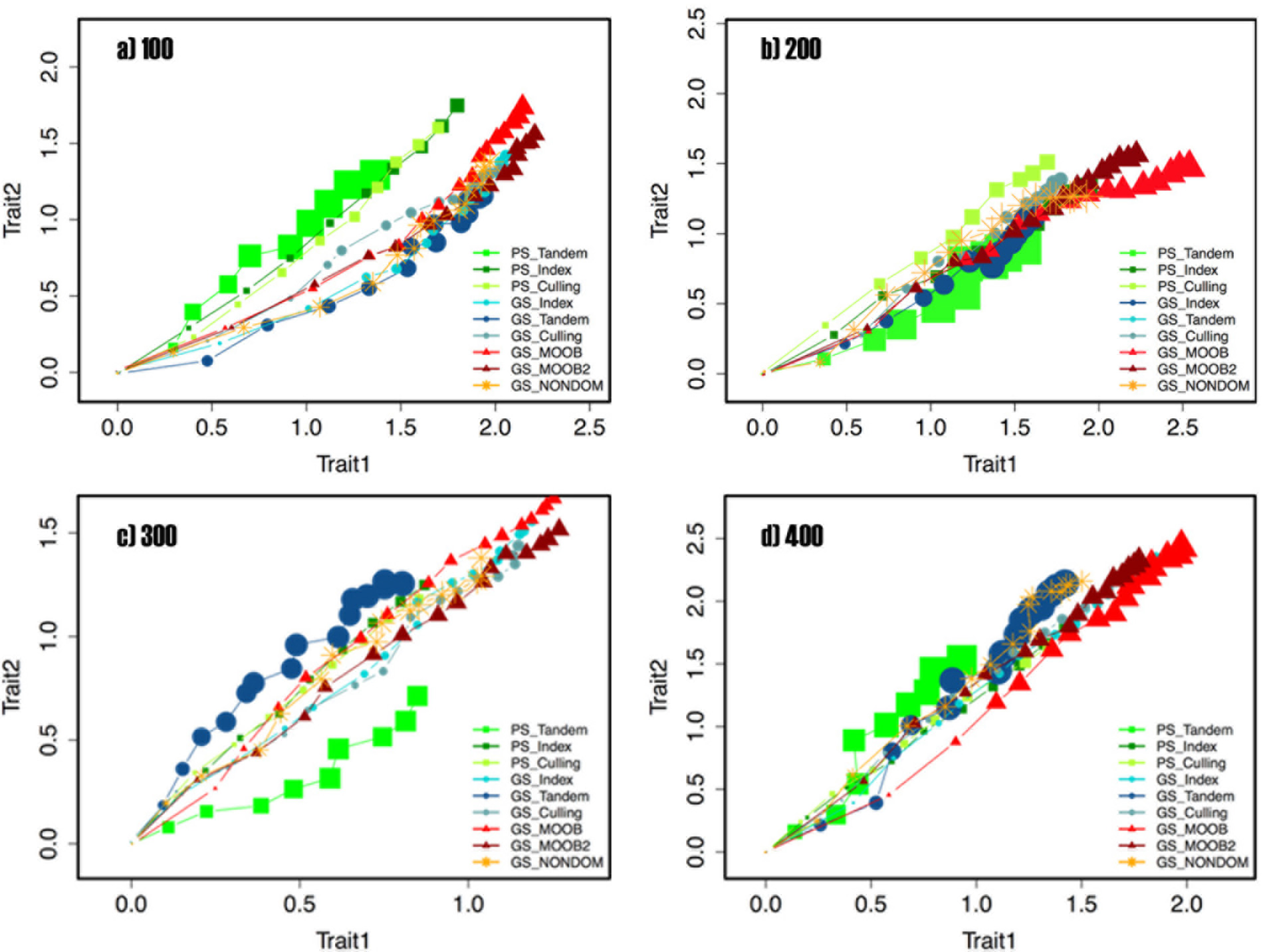
Simulations: the results from 30 replications of 16 rounds GS and 10 rounds PS with tandem, index selection and culling and 16 rounds with three multi-trait breeding methods. Breeding population sizes 100, 200, 300, and 400.

The simulation results indicate that the multi-objective optimized breeding approach is more efficient than classical methods. For all the population size, the multi-objective optimized approach performed more efficienty than any other method. Selection of solutions on the frontier during these simulations were done using a weighted distance to the ideal solution (which was taken as the optimal values for the three statistics within the solutions on the frontier). The weights for populations of size 100 and 200 were fixed at .95, .025, and .025 corresponding to the measures of inbreeding, gain in trait one and gain in trait 2. We have increased the intensity of selection for populations of size 300 and 400 by changing these weights to .9, .05, and .05. Note that, these values will be population specific and were here fixed only for the purposes of computer simulation of many cycles of breeding as described above. Decision support tools described previously should be utilized when applying the multi-objective optimized breeding methods.

## 5 DISCUSSION

A major task of breeders is to increase the frequency of favorable alleles of quantitative traits, controlled by a large number of genes. After choice of germplasm, a breeder uses some type of cyclical selection program to maximize the genetic improvement of desired traits. The important aspect is that superior materials selected for the breeding population should be recombined to obtain a new breeding population. It is important to incorporate recurrent selection into classical breeding programs [Hallauer and Carena Filho].

Most breeding programs are concerned with simultaneous improvement of several traits. For example, although yield is usually the primary trait of interest for most crops; maturity, standability, grain quality, stalk quality, abiotic and biotic stress tolerance, etc. are also economically important traits. Simultaneous selection for several traits is necessary if recurrent selection methods are used. Selection that emphasizes only one trait can be detrimental to the overall agronomic performance of the germplasm [Hallauer and Carena Filho].

In this article, we have described the use of multi-objective optimization techniques and concepts to breeding. We have demonstrated these with two real datasets and with simulation studies. We believe that the main promise of these techniques is that they provide the breeders with decision support tools that allow optimal exploitation of the breeding material. As contrasted to the index selection methods which require a-priori decision on economic weights, the multi-objective optimized breeding methodology involves selection of a ’good’ solution among the optimal compromise solutions with the aid of decision support tools such as the plots of frontier surface, or by using the ideal solution concept, visualizations using SOMs.

## Funding information

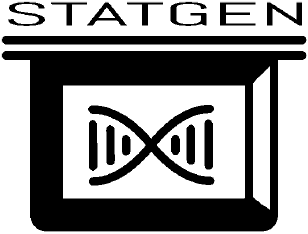

## CONFLICT OF INTEREST

Authors claim no conflict of interest.

## AUTHOR CONTRIBUTIONS

DA: Conception or design of the work, statistics, data preparation, programs, simulations, graphs, tables, drafting the article, critical revision of the article.

JS: Data preparation, drafting the article, graphs, tables, critical revision of the article.

## METHODS

### Multi-objective optimization techniques

In general, it is not possible to find an analytic expression of the line or surface that contains the Pareto optimal solutions. Several techniques are used to find representative points on the Pareto frontier:

#### Linear combination

If a solution *x* to the general multi-objective problem is non inferior, then there exist *w*_*l*_ = 0, *l* = 1,2,…,*k* (*w*_*r*_ is strictly positive for some *r* = 1,2,…, *k*), and *λ*_*i*_ = 0, *l* = 1,2,…,*m*, such that:

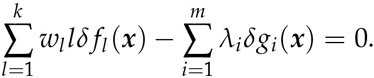

This condition is named Kuhn-Tucker Condition (KTC) and is necessary for a non inferior solution. When all of the *f*_*l*_(*x*) are concave and *x* belongs to a convex set, they are sufficient as well. Since KTC is sufficient for non-inferiority, non inferior solutions might be found by solving a scalar optimization problem in which the objective function is a weighted sum of the components of the original vector-valued function. That is, the solution to the problem: 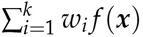, where *w*_*i*_ ≥ 0 for all *i* and strictly positive for at least one objective, is usually non inferior. Then on inferior set and the set of non inferior solutions can be generated by varying the weights *w*_*i*_ in the objective function.

The reduction of the problem to a single-objective function means to make all alternatives comparable with a preference framework that becomes a total order. Hence *w*_*i*_ values choice is very important to achieve the final decision and, for this reason, value choice is made by the decision maker. However the decision maker, in order to choose the coefficients, must have a clear perception of how this choice effects all the functions with respect to each other.

The main advantages of this method are its simplicity (in implementation and use) and its efficiency (computationally speaking). Its main disadvantage is the difficulty to determine the appropriate weight coefficients to be used when enough information about the problem is not available (this is an important concern, particularly in real-world applications). Also, a proper scaling of the objectives requires a considerable amount of extra knowledge about the problem. To obtain this information could be a very expensive process. A more serious drawback of this approach, is that it cannot be used to generate certain portions of the Pareto front when the conditions of KTC are not satisfied, regardless of the weights combination used. Nevertheless, aggregating functions could be very useful to get a preliminary sketch of the Pareto front of a certain problem or to provide prior information to be exploited by another approach.

Other scalarization methods include *L*_*p*_-norm, Chebyshev and the single-objective product formulation. For each of these scalarizations, a characterization of the Pareto set can be obtained by varying the scalarization parameters and solving many single-objective optimization problems. Ideally, the points returned by the scalarized problems should be sufficiently spread out in the efficient frontier.

#### The *∊*-constraint method

Besides the linear combination approach, the *∊*-constraint method is probably the best known technique to solve multi-objective optimization problems. There is no aggregation of criteria, instead only one of the original objectives is minimized while the others are transformed to constraints. The idea was introduced by Haimes [16]. Through this approach among *p* objective function only one is kept as such, the other *p*-1 are transformed in constraints fixing threshold values *∊*_*k*_ (with *k* = 1,…,*p*, *k* ≠ *j*) over them (if functions must be minimized). Therefore the problem: *min*F(*x*) is substituted by the *∊*-constraint problem: *min f*_*j*_(*x*) *f*_*k*_(*x*) ≤ *∊*_*k*_, *k* = 1,…,*p*, *k* ≠ *j*. The main disadvantage of this approach is its (potentially high) computational cost, also due to the preliminary individuation of *∊*_*i*_ values.

#### Finding non-dominated solutions

Several algorithms exist for finding the non-dominated set in from a larger set. Popular Kung algorithm ([1]) involves first sorting the population in descending order in accordance to first objective function. Afterwards, the population is recursively partitioned as top (T) and bottom (B) sub-populations. As top half (T) is better in objective in comparison to bottom half (B) in first objective, so we check the bottom half for domination with top half. The solution of B which are not dominated by solutions of T are merged with members of T to form merged population M. Another algorithm is the Jun Du Algorithm ([2]).

#### Other multi-trait breeding approaches

Figures S1 and S2 display the individuals that would be identified by the classical multi-trait breeding schemes culling, tandem and index selection for the wheat and barley datasets. These can be contrasted with the Figures 1 a and b and also with the Figures3 and S5. Note that among the classical methods index selection will give the closest results to the multi-objective optimized breeding methods.

**Fig. S1.**
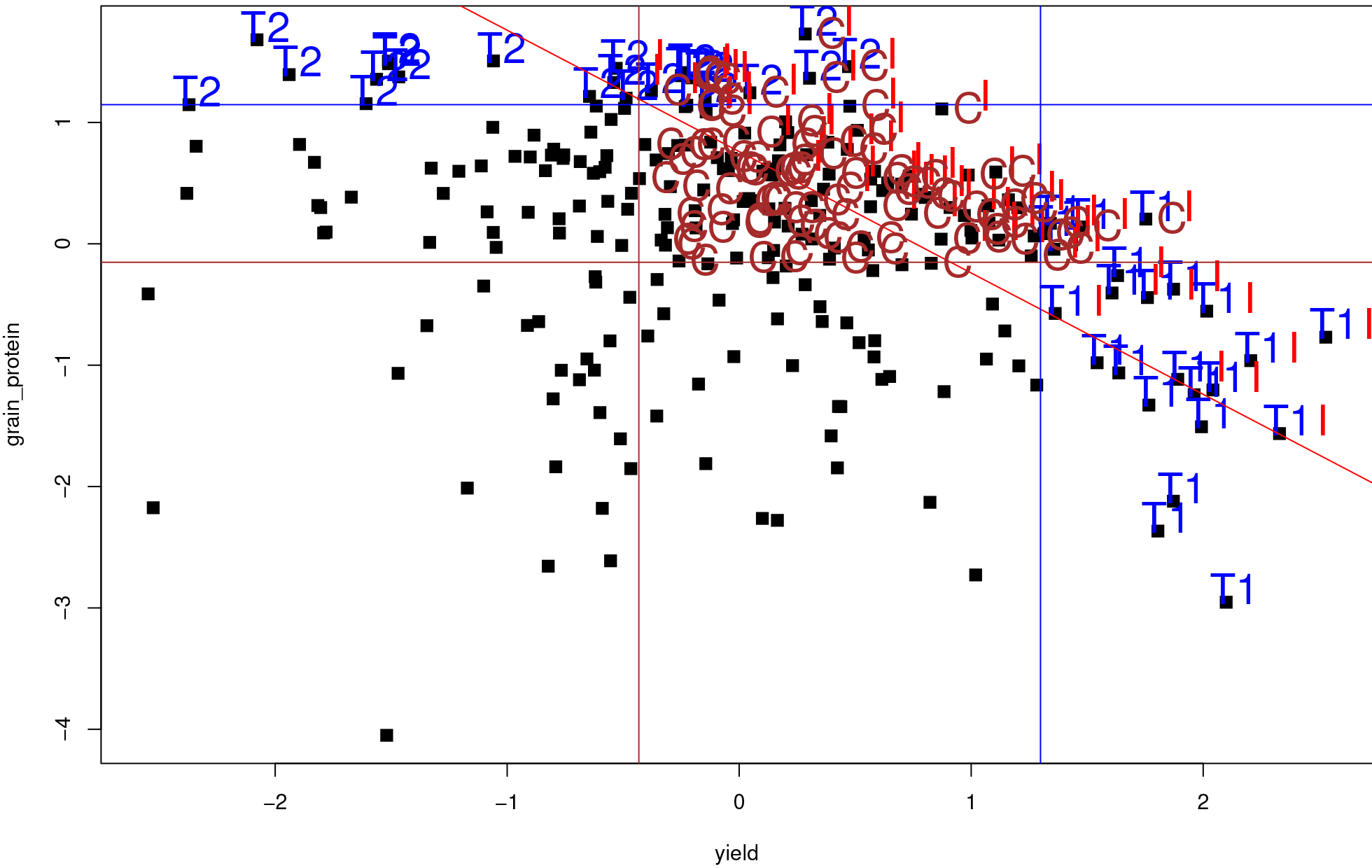
Wheat data: Other multi-trait breeding approaches

**Fig. S2.**
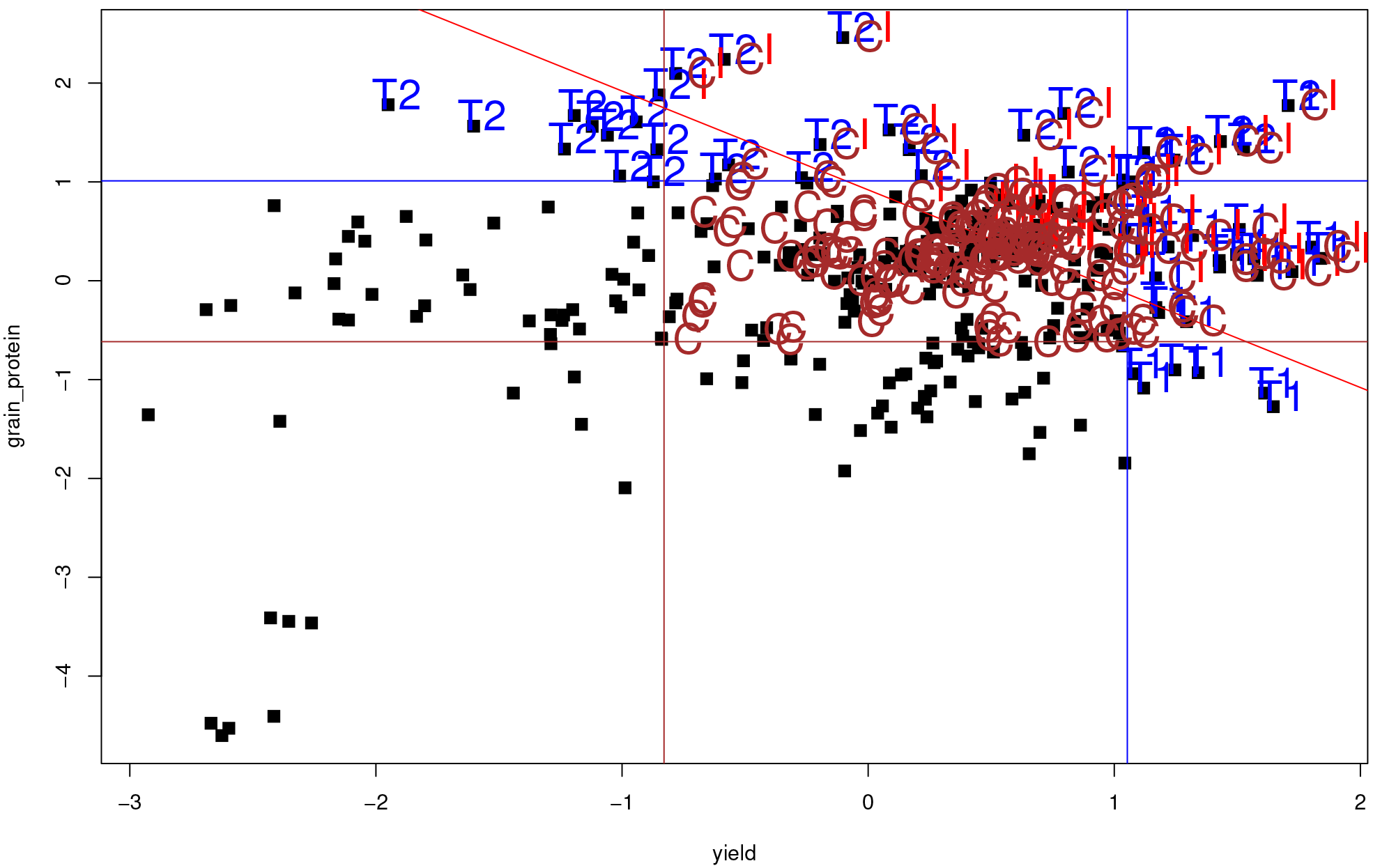
Barley data: Other multi-trait breeding approaches

## ADDITIONAL FIGURES

**Fig. S3.**
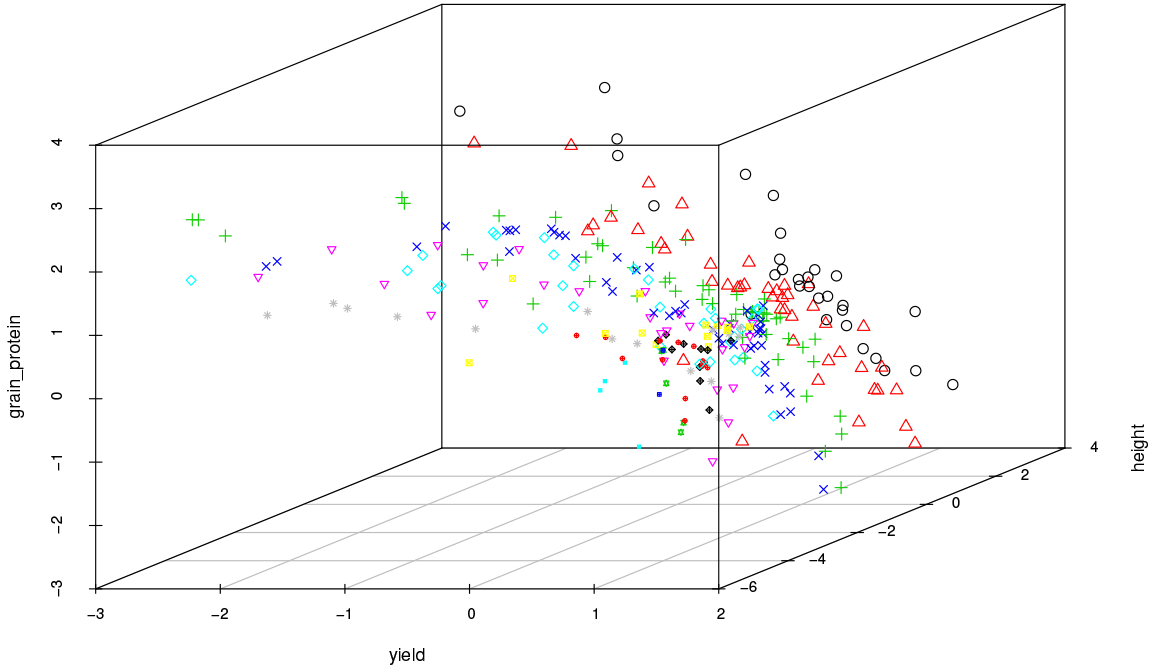
This figure represents the non-dominance ordering of the individuals in the barley dataset for three traits. Barley data: Dominance ordering based on three traits, 13 levels of dominance. The GEBVs for height, grain yield and protein from barley data are plotted with the dominance ordering of these individuals indicated by the lines of different color.

Figure S4 displays the parental contribution proportions obtained by non-dominance counts for 100 lines with the highest proportions for the improving yield and protein in the four environments.

**Fig. S4.**
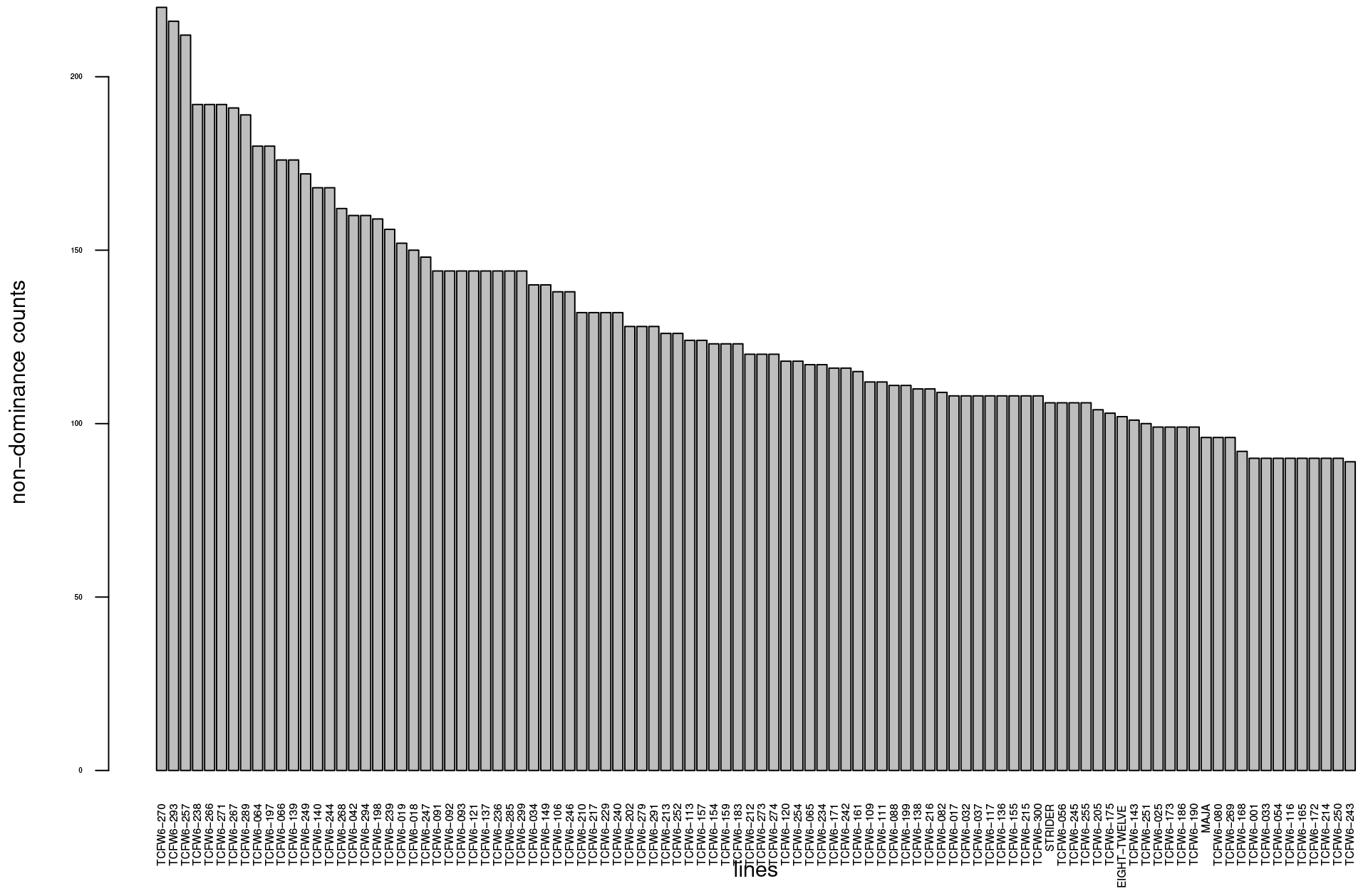
Barley data: parental contribution proportions obtained by non-dominance counts for 100 lines with the highest proportions for the improving yield in four environments.

**Fig. S5.**
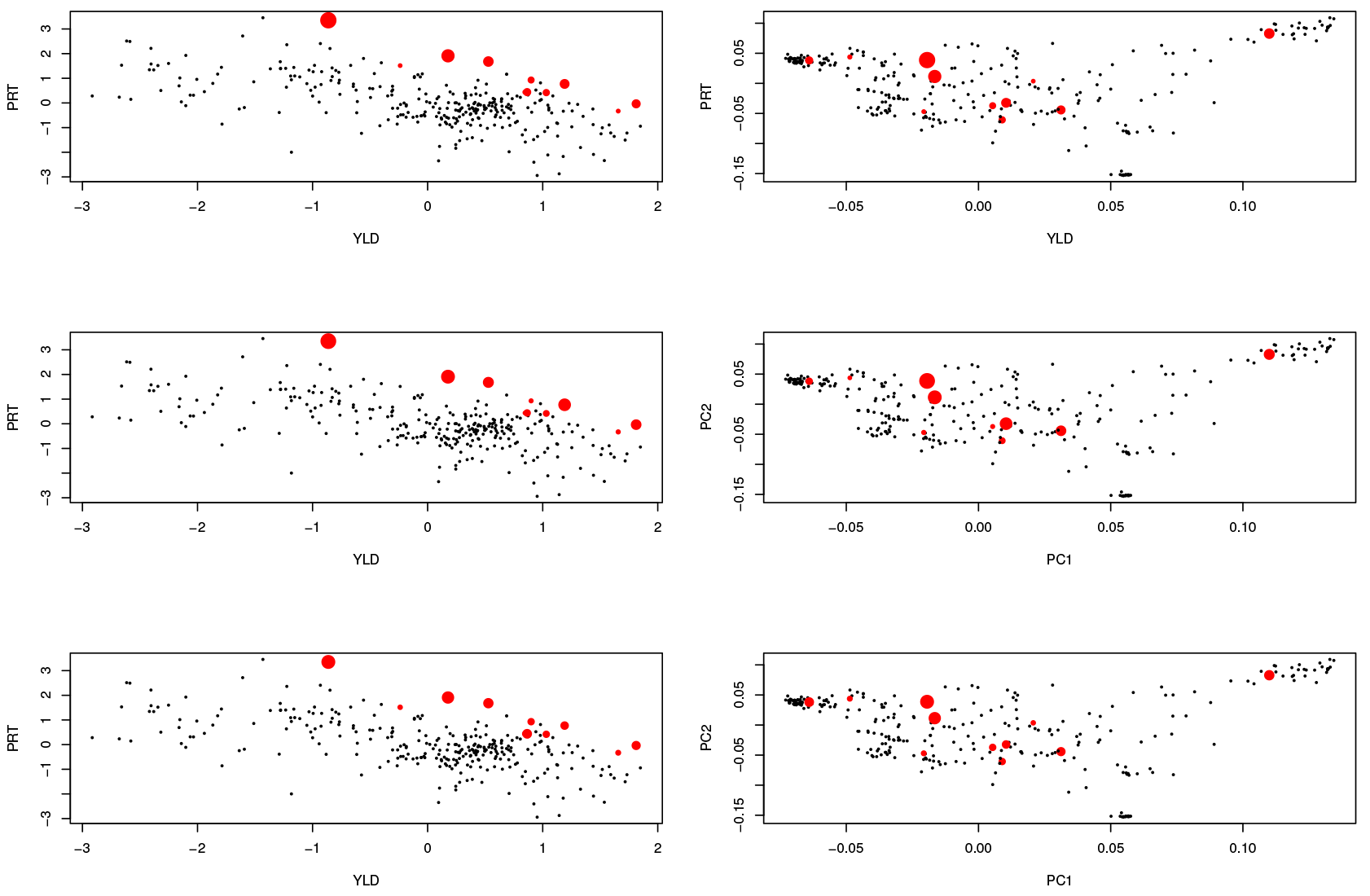
Three ’good’ solutions on the barley frontier curve obtained from figure 2. Red points indicated the individuals that have non-zero parental proportions. The size of the points are proportional to the magnitude of the parental contributions. The figures on the right side, represent the same information but on the first two principal components of the genotyping marker space.)

**Fig. S6.**
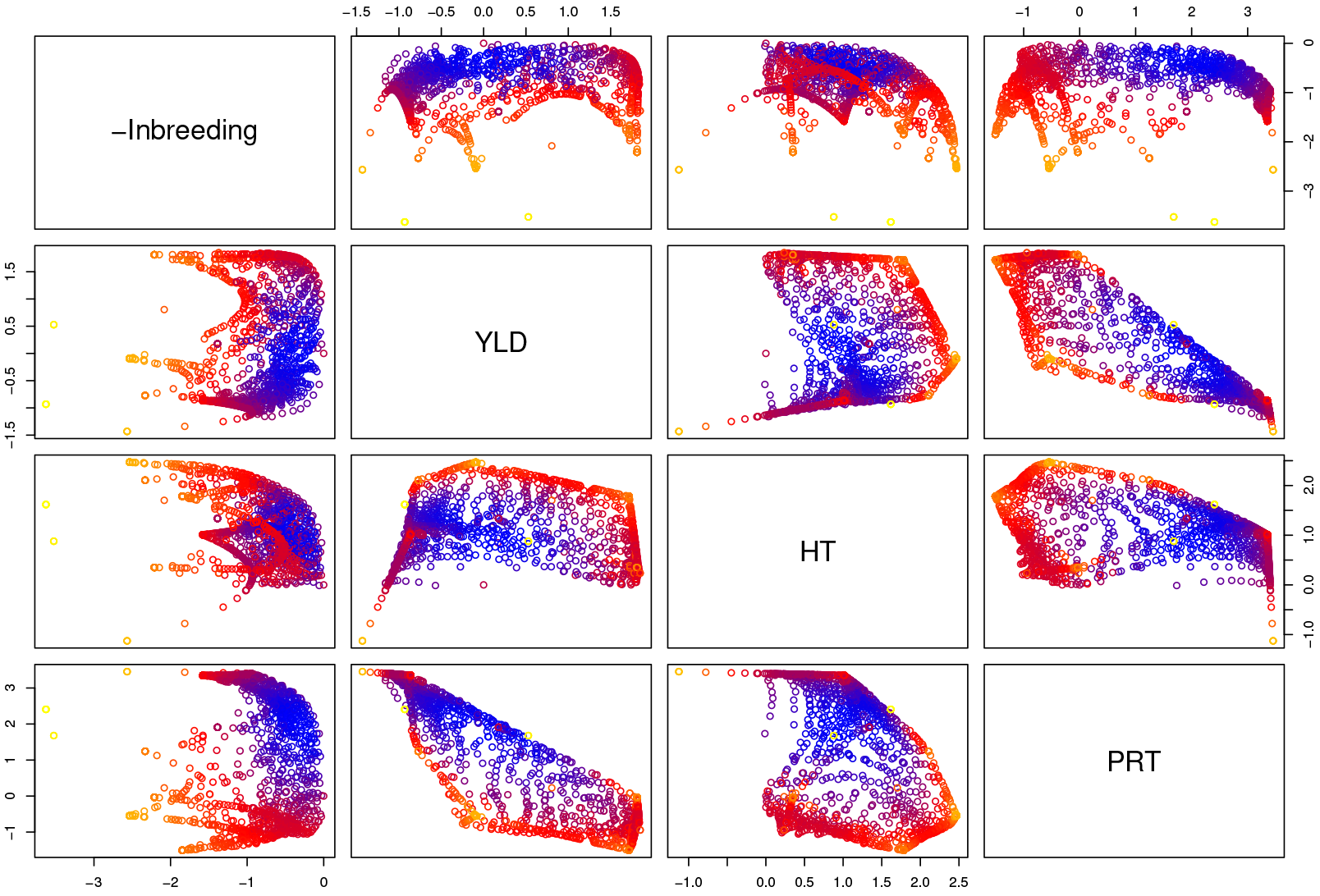
Barley data: Four dimensional Pareto front represented in two dimensional plots. The dimensions correspond to negative of inbreeding, and average gains based on genomically estimated breeding values for yield, height and protein content. Each point on the scatterplots correspond to a Pareto optimal solution for parental contributions. Blueness of the points measure the closeness to the ideal solution.

**Fig. S7.**
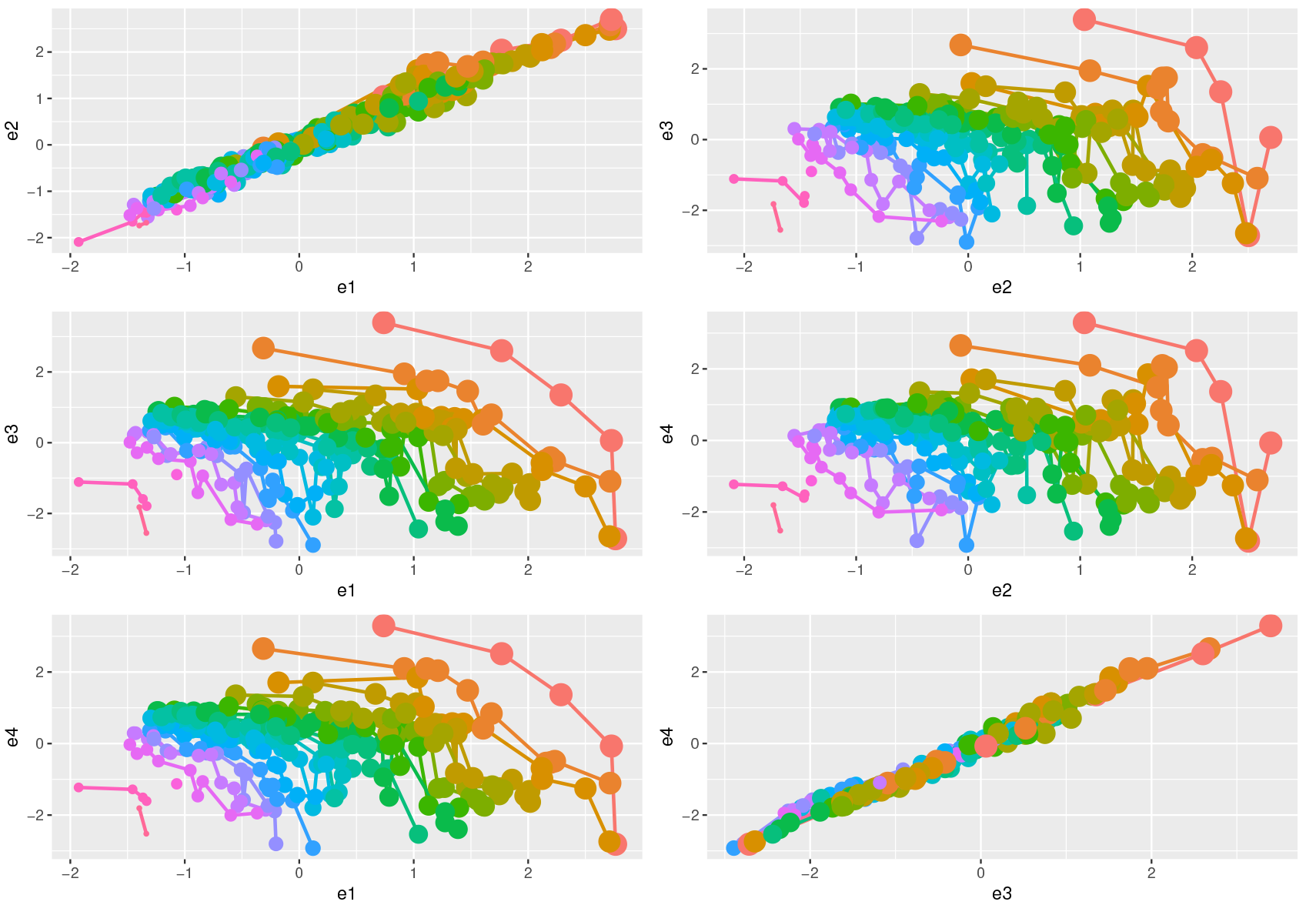
Barley data: Dominance ordering based on protein content in 4 environments‥ The environment specific GEBVs for protein content for barley data are plotted with the dominance ordering of these individuals indicated by the lines of different color.

**Fig. S8.**
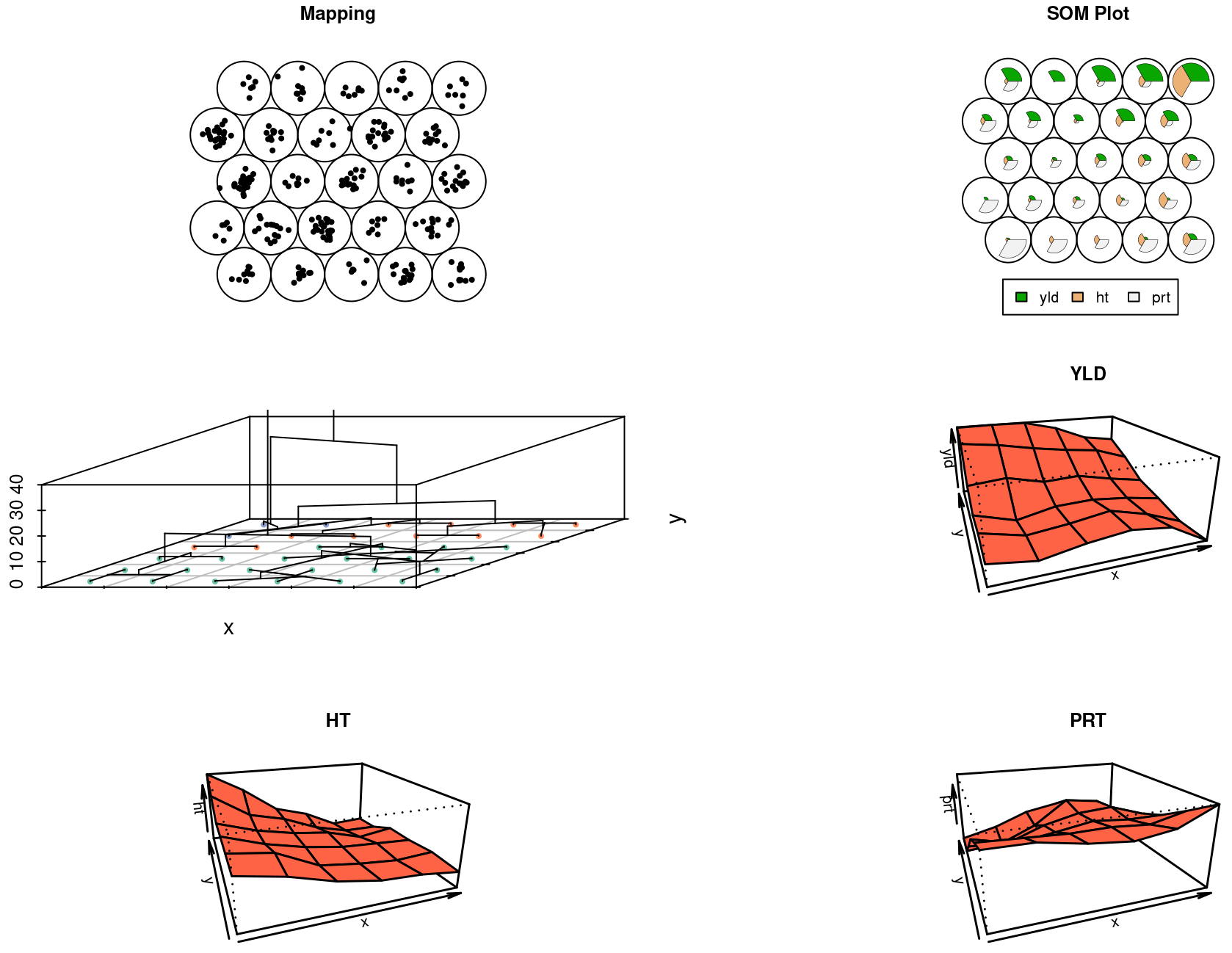
Barley data: SOM plot for barley data for GEBVs over three traits, yield, protein content and height.

**Fig. S9.**
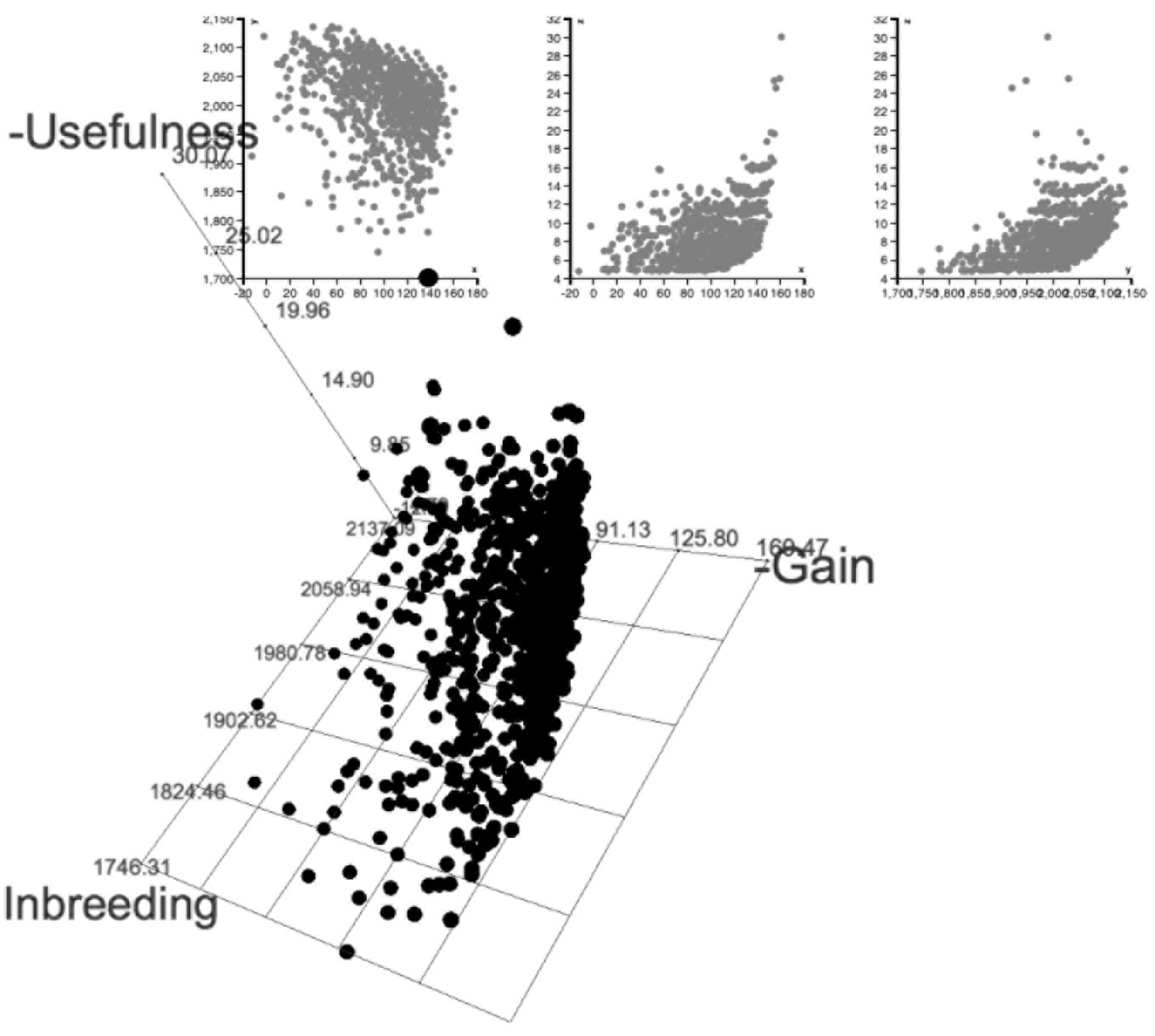
Simulated data: Gain, Usefulness, Coancestory: Genomic mating approach is also a multiobjective optimized breeding approach. The frontier surface in the figure represents the tradeoff between the measures of gain, usefulness, coancestory for pareto optimal mating plans.

**Fig. S10.**
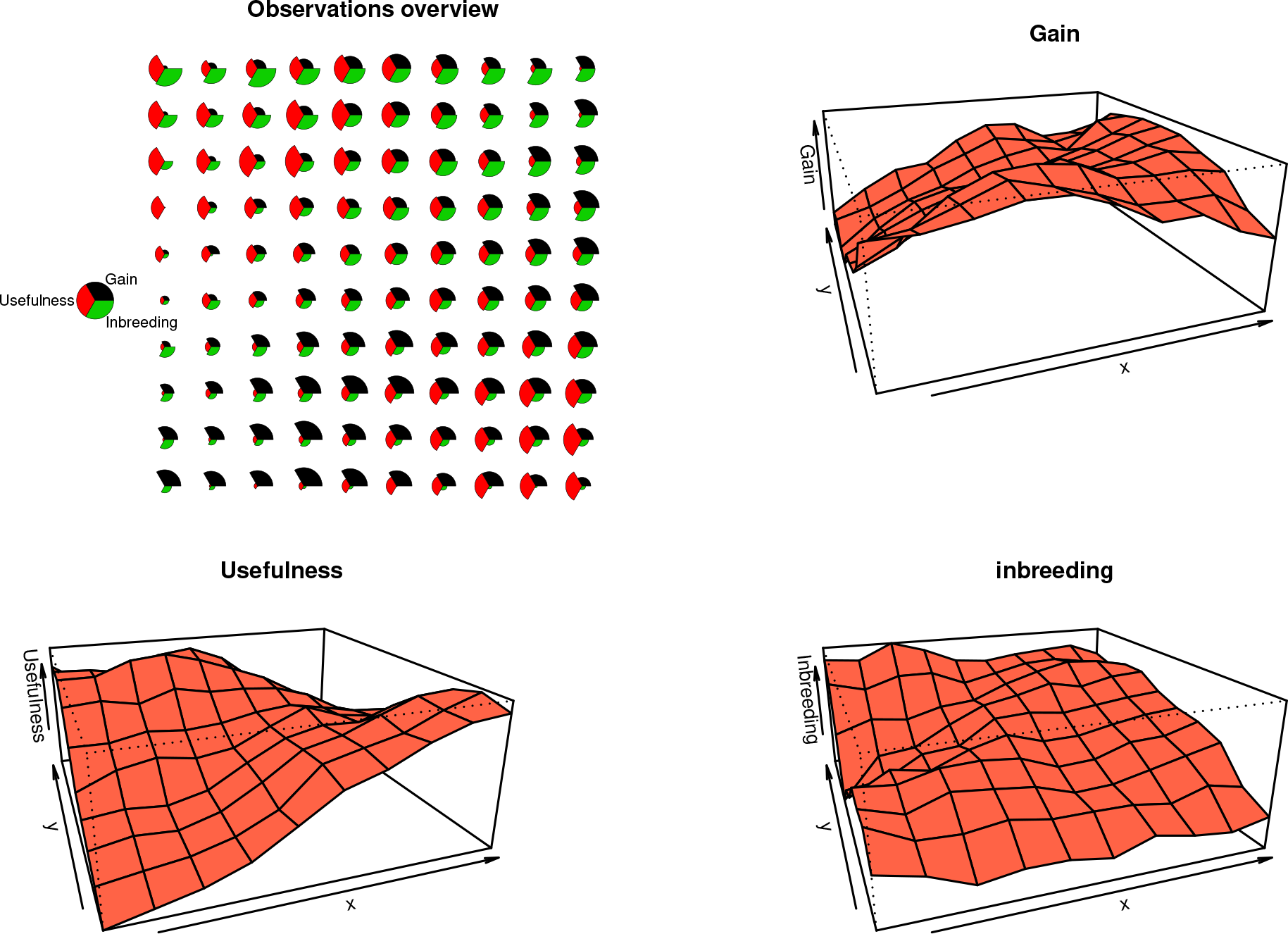
Simulated data: Gain, Usefulness, Coancestory: The top left figure is a representation of all the solutions on the Pareto frontier in Figure S9 on a two dimensional SOM. The remaining graphs are showing how each measure behaves individually on the two SOM features.

## CONFLICT OF INTEREST

Authors claim no conflict of interest.

## AUTHOR CONTRIBUTIONS

DA: Conception or design of the work, programs, simulations, graphs, drafting the article, critical revision of the article.

JS: Graphs, drafting the article, critical revision of the article.

## REFERENCES

Acquaah, G. (2009) Principles of plant genetics and breeding. John Wiley & Sons.

Agrawal, G., Bloebaum, C. and Lewis, K. (2005) Intuitive design selection using visualized n-dimensional pareto frontier. In 1st AIAA Multidisciplinary Design Optimization Specialist Conference.

Aiking, H. (2011) Future protein supply. Trends in Food Science & Technology, 22, 112–120.

Akdemir, D. and Sánchez, J. I. (2016) Efficient breeding by genomic mating. Frontiers in genetics, 7, 210.

Allard, R. W. (1999) Principles of plant breeding. John Wiley & Sons.

Bernardo, R. (2002) Breeding for quantitative traits in plants, vol. 1. Stemma Press.

Bernardo, R. and Charcosset, A. (2006) Usefulness of gene information in marker-assisted recurrent selection: a simulation appraisal. Crop Science, 46, 614–621.

Bernardo, R. and Yu, J. (2007) Prospects for genomewide selection for quantitative traits in maize. Crop Science, 47, 10821090.

Brisbane, J. and Gibson, J. (1995) Balancing selection response and rate of inbreeding by including genetic relationships in selection decisions. Theoretical and Applied Genetics, 91, 421–431.

Burgess, J. C. and West, D. (1993) Selection for grain yield following selection for ear height in maize. Crop science, 33, 679–682.

Casadebaig, P., Mestries, E. and Debaeke, P. (2016) A model-based approach to assist variety evaluation in sunflower crop. European Journal of Agronomy, 81, 92–105.

Charcosset, A. and Hospital, F. (1997) Marker-assisted introgression of quantitative trait loci. Genetics, 147, 1469–1485.

Clark, S. A., Kinghorn, B. P., Hickey, J. M. and van der Werf, J. H. (2013) The effect of genomic information on optimal contribution selection in livestock breeding programs. Genetics Selection Evolution, 45, 1.

Cockerham, C. C. (1963) Estimation of genetic variances. Statistical genetics and plant breeding, 982, 53–94.

Coello, C. C. (2006) Evolutionary multi-objective optimization: a historical view of the field. IEEE computational intelligence magazine, 1, 28–36.

Cooper, M. and DeLacy, I. (1994) Relationships among analytical methods used to study genotypic variation and genotype-by-environment interaction in plant breeding multi-environment experiments. TAG Theoretical and Applied Genetics, 88, 561–572.

Cooper, M., Woodruff, D., Phillips, I., Basford, K. and Gilmour, A. (2001) Genotype-by-management interactions for grain yield and grain protein concentration of wheat. Field Crops Research, 69, 47–67.

Crossa, J., Burgueño, J., Cornelius, P. L., McLaren, G., Trethowan, R. and Krishnamachari, A. (2006) Modeling genotype× environment interaction using additive genetic covariances of relatives for predicting breeding values of wheat genotypes. Crop science, 46,1722–1733.

Daetwyler, H. D., Villanueva, B., Bijma, P. and Woolliams, J. A. (2007) Inbreeding in genome-wide selection. Journal of Animal Breeding and Genetics, 124, 369–376.

Deb, K. (2001) Multi-objective optimization using evolutionary algorithms, vol. 16. John Wiley & Sons.

DePauw, R., Knox, R., Clarke, F., Wang, H., Fernandez, M., Clarke, J. and McCaig, T. (2007) Shifting undesirable correlations. Euphytica, 157, 409–415.

Dudley, J. and Moll, R. (1969) Interpretation and use of estimates of heritability and genetic variances in plant breeding. Crop science, 9, 257–262.

Falconer, D. S., Mackay, T. F. and Frankham, R. (1996) Introduction to quantitative genetics (4th edn). Trends in Genetics, 12, 280.

Fernández, J., Toro, M. and Caballero, A. (2001) Practical implementation of optimal management strategies in conservation programmes: a mate selection method. Animal Biodiversity and Conservation, 24, 17–24.

Ghanem, M. E., Marrou, H. and Sinclair, T. R. (2015) Physiological phenotyping of plants for crop improvement. Trends in Plant Science, 20, 139–144.

Gianola, D. and Fernando, R. L. (1986) Bayesian methods in animal breeding theory. Journal of Animal Science, 63, 217–244.

Goddard, M. (2009) Genomic selection: prediction of accuracy and maximisation of long term response. Genetics, 136, 245257.

Gouache, D., Bogard, M., Pegard, M.,Thepot, S., Garcia, C., Hourcade, D., Paux, E., Oury, F.-X., Rousset, M., Deswarte, J.-C. et al. (2017) Bridging the gap between ideotype and genotype: Challenges and prospects for modelling as exemplified by the case of adapting wheat (triticum aestivum l.) phenology to climate change in france. Field Crops Research, 202, 108–121.

Groos, C., Robert, N., Bervas, E. and Charmet, G. (2003) Genetic analysis of grain protein-content, grain yield and thousand-kernel weight in bread wheat. TAG Theoretical and Applied Genetics, 106, 1032–1040.

Hallauer, A. and Carena Filho, M. () Jbm (2010) quantitative genetics in maize breeding, handbook of plant breeding.

Hallauer, A. R. and Miranda, J. (1987) Quantitative genetics in maize breeding. Iowa State University Pres. Ames.

Hazel, L. and Lush, J. L. (1942) The efficiency of three methods of selection. Journal of Heredity, 33, 393–399.

Hazel, L. N. (1943) The genetic basis for constructing selection indexes. Genetics, 28, 476–490.

Heffner, E. L., Sorrells, M. E. and Jannink, J.-L. (2009) Genomic selection for crop improvement. Crop Science, 49, 1–12.

Henderson, C. (1984) Applications of linear models in animal breeding (University of Guelph, Guelph, ON, Canada). Elsevier.

Holland, J. B., Nyquist, W. E. and Cervantes-Martínez, C. T. (2003) Estimating and interpreting heritability for plant breeding: an update. Plant breeding reviews, 22, 9–112.

Jannink, J.-L., Lorenz, A. J. and Iwata, H. (2010) Genomic selection in plant breeding: from theory to practice. Briefings in functional genomics, elq001.

Kibite, S. and Evans, L. (1984) Causes of negative correlations between grain yield and grain protein concentration in common wheat. Euphytica 33, 801–810.

Kinghorn, B. and Shepherd, R. (1999) Mate selection for the tactical implementation of breeding programs. Association Advancement Animal Breeding Genetics, 13, 130–133.

Kohonen, T. (1981) Automatic formation of topologicalmaps of patterns in a self-organizing system. Scand. Conf. on Image Analysis, 214–220.

Kohonen, T. (1998) The self-organizing map. Neurocomputing, 21, 1–6.

Konak, A., Coit, D. W. and Smith, A. E. (2006) Multi-objective optimization using genetic algorithms: A tutorial. Reliability Engineering & System Safety, 91, 992–1007.

Lande, R. and Thompson, R. (1990) Efficiency of marker-assisted selection in the improvement of quantitative traits. Genetics, 124, 743–756.

Lander, E. S., Linton, L. M., Birren, B., Nusbaum, C., Zody, M. C., Baldwin, J., Devon, K., Dewar, K., Doyle, M., FitzHugh, W. et al. (2001) Initial sequencing and analysis of the human genome. Nature 409, 860–921.

Legarra, A., Aguilar, I. and Misztal, I. (2009) A relationship matrix including full pedigree and genomic information. Journal of dairy science, 92, 4656–4663.

Leutenegger, A.-L., Prum, B., Génin, E., Verny, C., Lemainque, A., Clerget-Darpoux, F. and Thompson, E. A. (2003) Estimation of the inbreeding coefficient through use of genomic data. The American Journal of Human Genetics, 73, 516–523.

Lynch, M., Walsh, B. et al. (1998) Genetics and analysis of quantitative traits, vol. 1. Sinauer Sunderland, MA.

Margulies, M., Egholm, M., Altman, W. E., Attiya, S., Bader, J. S., Bemben, L. A., Berka, J., Braverman, M. S., Chen, Y.-J., Chen, Z. et al. (2005) Genome sequencing in microfabricated high-density picolitre reactors. Nature, 437, 376–380.

Martre, P., He, J., Le Gouis, J. and Semenov, M. A. (2015a) In silico system analysis of physiological traits determining grain yield and protein concentration for wheat as influenced by climate and crop management. Journal of experimental botany, 66, 3581–3598.

Martre, P., Quilot-Turion, B., Luquet, D., Memmah, M.-M. O.-S., Chenu, K. and Debaeke, P. (2015b) Model-assisted phenotyping and ideotype design. Crop physiology: applications for genetic improvement and agronomy, 349–373.

Metzker, M. L. (2010) Sequencing technologies-the next generation. Nature reviews genetics, 11, 31–46.

Meuwissen, T. (1997) Maximizing the response of selection with a predefined rate of inbreeding. Journal of animal science, 75, 934–940.

Meuwissen, T. H. E., Hayes, B. J. and Goddard, M. E. (2001) Prediction of total genetic value using genome-wide dense marker maps. Genetics, 157, 1819–1829.

Moose, S. P. and Mumm, R. H. (2008) Molecular plant breeding as the foundation for 21st century crop improvement. Plant physiology, 147, 969–977.

Obayashi, S. and Sasaki, D. (2003) Visualization and data mining of pareto solutions using self-organizing map. In EMO, 796809. Springer.

Oury, F.-X. and Godin, C. (2007) Yield and grain protein concentration in bread wheat: how to use the negative relationship between the two characters to identify favourable genotypes? Euphytica, 157, 45–57.

Picheny, V., Casadebaig, P., Trépos, R., Faivre, R., Da Silva, D., Vincourt, P. and Costes, E. (2017) Using numerical plant models and phenotypic correlation space to design achievable ideotypes. Plant, Cell & Environment.

Piepho, H., Möhring, J., Melchinger, A. and Büchse, A. (2008) Blup for phenotypic selection in plant breeding and variety testing. Euphytica, 161, 209–228.

Pryce, J., Hayes, B. and Goddard, M. (2012) Novel strategies to minimize progeny inbreeding while maximizing genetic gain using genomic information. Journal of Dairy Science, 95, 377–388.

Quaas, R. (1988) Additive genetic model with groups and relationships. Journal of Dairy Science, 71, 1338–1345.

Rharrabti, Y., Villegas, D., García del Moral, L., Aparicio, N., Elhani, S. and Royo, C. (2001) Environmental and genetic determination of protein content and grain yield in durum wheat under mediterranean conditions. Plant Breeding, 120, 381–388.

Rötter, R., Tao, F., Höhn, J. and Palosuo, T. (2015) Use of crop simulation modelling to aid ideotype design of future cereal cultivars. Journal of experimental botany, 66, 3463–3476.

Schierenbeck, S., Pimentel, E., Tietze, M., Körte, J., Reents, R., Reinhardt, F., Simianer, H. and König, S. (2011) Controlling inbreeding and maximizing genetic gain using semi-definite programming with pedigree-based and genomic relationships. Journal of dairy science, 94, 6143–6152.

Shepherd, R. and Kinghorn, B. (1998) A tactical approach to the design of crossbreeding programs. In Proceedings of the Sixth World Congress on Genetics Applied to Livestock Production: 11-16 January; Armidale, vol. 25, 431–438.

Shewry, P. R. (2007) Improving the protein content and composition of cereal grain. Journal of Cereal Science, 46, 239–250.

Simmonds, N. W. (1995) The relation between yield and protein in cereal grain. Journal of the Science of Food and Agriculture, 67, 309–315.

Smith, H. F. (1936) A discriminant function for plant selection. Annals of Human Genetics, 7, 240–250.

Sonesson, A. K., Woolliams, J. A. and Meuwissen, T. H. (2012) Genomic selection requires genomic control of inbreeding. Genetics Selection Evolution, 44, 1.

Sun, C., VanRaden, P., O’Connell, J., Weigel, K. and Gianola, D. (2013) Mating programs including genomic relationships and dominance effects. Journal of Dairy Science, 96, 8014–8023.

Tušar, T. and Filipič, B. (2015) Visualization of pareto front approximations in evolutionary multiobjective optimization: A critical review and the prosection method. IEEE Transactions on Evolutionary Computation, 19, 225–245.

de la Vega, A. (2012) Effect of the complexity of sunflower growing regions on the genetic progress achieved by breeding programs. Helia, 35, 113–122.

Wang, J. (2011) Coancestry: a program for simulating, estimating and analysing relatedness and inbreeding coefficients. Molecular ecology resources, 11, 141–145.

Williams, J. (1962) The evaluation of a selection index. Biometrics, 18, 375–393.

Wray, N. and Goddard, M. (1994) Moet breeding schemes for wool sheep 1. design alternatives. Animal Production, 59, 71–86.

Wright, S. (1921) Systems of mating. i. the biometric relations between parent and offspring. Genetics, 6, 111.

Zheng, B. S., Le Gouis, J., Daniel, D. and Brancourt-Hulmel, M. (2009) Optimal numbers of environments to assess slopes of joint regression for grain yield, grain protein yield and grain protein concentration under nitrogen constraint in winter wheat. Field Crops Research, 113, 187–196.

Zhong, S. and Jannink, J.-L. (2007) Using quantitative trait loci results to discriminate among crosses on the basis of their progeny mean and variance. Genetics, 177, 567–576.

Zio, E. and Bazzo, R. (2012) A comparison of methods for selecting preferred solutions in multiobjective decision making. Computational intelligence systems in industrial engineering, 23–43.

## REFERENCES

1. H.-T. Kung, F. Luccio, and F. P. Preparata, “On finding the maxima of a set of vectors,” Journal of the ACM (JACM) 22, 469–476 (1975).

2. J. Du and Z. Cai, “A sorting based algorithm for finding a non-dominated set in multi-objective optimization,” in “Natural Computation, 2007. ICNC 2007. Third International Conference on,”, vol. 4 (IEEE, 2007), vol. 4, pp. 436–440.

